# Assembly of tight junction belts by surface condensation and actin elongation

**DOI:** 10.1101/2023.06.24.546380

**Authors:** Daxiao Sun, Xueping Zhao, Tina Wiegand, Giacomo Bartolucci, Cecilie Martin-Lemaitre, Stephan W. Grill, Anthony A. Hyman, Christoph Weber, Alf Honigmann

## Abstract

Formation of biomolecular condensates via phase separation enables compartmentation of many cellular processes. However, how cells can control condensation at specific locations to create complex cellular structures remains poorly understood. Here, we investigated the mechanism of tight junction formation, which involves condensation of scaffold proteins at cell-cell contacts and elongation of the condensates into a belt around the cellular perimeter. Using cell biology, reconstitution, and thermodynamic theory, we discovered that cells use surface phase transitions to control local condensation at the membrane far below bulk saturation. Surface condensation of junctional ZO-scaffold proteins is mediated by receptor binding and regulated by the receptor’s oligomerization state. Functionally, ZO surface condensation is directly coupled to actin polymerization and bundling, which drives elongation of receptor-ZO-actin condensates similar to tight junction belt formation in cells. We conclude that surface phase transitions provide a robust mechanism to control the position and shape of protein condensates.

**One-Sentence Summary:** Local surface binding of cytosolic scaffold proteins provides spatial control of protein condensation to assemble adhesion junctions.

## Introduction

Tight junctions are adhesion complexes that control the paracellular flux of solutes across tissues (*1-3*). The super-molecular structure of tight junctions is constructed from adhesion receptors of the claudin (CLDN) family, which form intercellular adhesion strands that act as diffusion barriers, and Zonula occludens (ZO) scaffold proteins which connect the receptors to the cytoskeleton (*1*). Assembly of junctions is initiated by condensation of cytosolic scaffold ZO proteins at cell-cell contact sites that over time elongate and fuse around the apical cell perimeters into a continuous belt which seals the tissue (*4-6*). How cells spatially control condensation at cell-cell contacts and how the condensates are reshaped into closed belts has remained unclear.

The formation of protein condensates in the cytoplasm (*7-9*) and nucleoplasm (*10, 11*) can be governed by thermodynamics (*12*). Formation of condensates via phase separation requires protein concentrations above the bulk saturation concentration (*13*). However, on membrane surfaces condensate formation has been observed even far below bulk saturation (*14-16*), which suggests distinct phase transitions at the surface in comparison to the bulk. Surface phase transitions (*17*) including the prewetting transition have been originally predicted by Cahn (*18*) and have been found to be close to the saturation concentration in experimental polymeric systems (*19-21*). However, a recent theoretical study found that phase transitions can occur far below saturation if molecules can bind to the surface (*22*). Indeed, a prewetting surface phase transition was proposed as a mechanism for condensation of transcription factors binding to DNA surfaces (*23, 24*). How surface phase transitions are spatially controlled on biological membranes is not well understood.

In this work, we established how cells can induce a surface phase transition specifically at cell-cell contact regions and how subsequent interactions of surface condensates with the cytoskeleton drive its active remodeling into an elongated and functional tight junction belt.

## Results

To explore the mechanism of formation and maturation of junctional condensates into a belt structure, we reconstituted the tight junction assembly pathway on model membranes from purified components focusing on the intracellular components of the junction (Figure 1A), which comprise the cytoplasmic tails of adhesion receptors (CLDN2, JAM-A, CDH1), the scaffold proteins (ZO1-3), adapter proteins (CGN, AFDN, PAR3) and the cytoskeleton (Actin) (Fig. S1, A and B). Initially, we aimed at characterizing the interactions of the main scaffold protein ZO1 to the strand forming receptor CLDN2 (Fig. 1A). We prepared fluorescently labeled C-terminal tails of CLDN2 containing the PDZ-binding motif for ZO1 and an N-terminal His-tag that allowed its attachment to Ni^2+^-containing supported lipid bilayer (*14, 15*). During cell-cell adhesion CLDN receptors are known to oligomerize (*25, 26*). To mimic different oligomerization states of the CLDN2 receptor, we attached GCN4 and ySMF domains that assemble into stable tetramers or 14-mers, respectively (*27, 28*). This setup allowed us to systematically titrate receptor surface density and ZO1 bulk concentration to characterize binding and condensate formation.

**Fig. 1.**
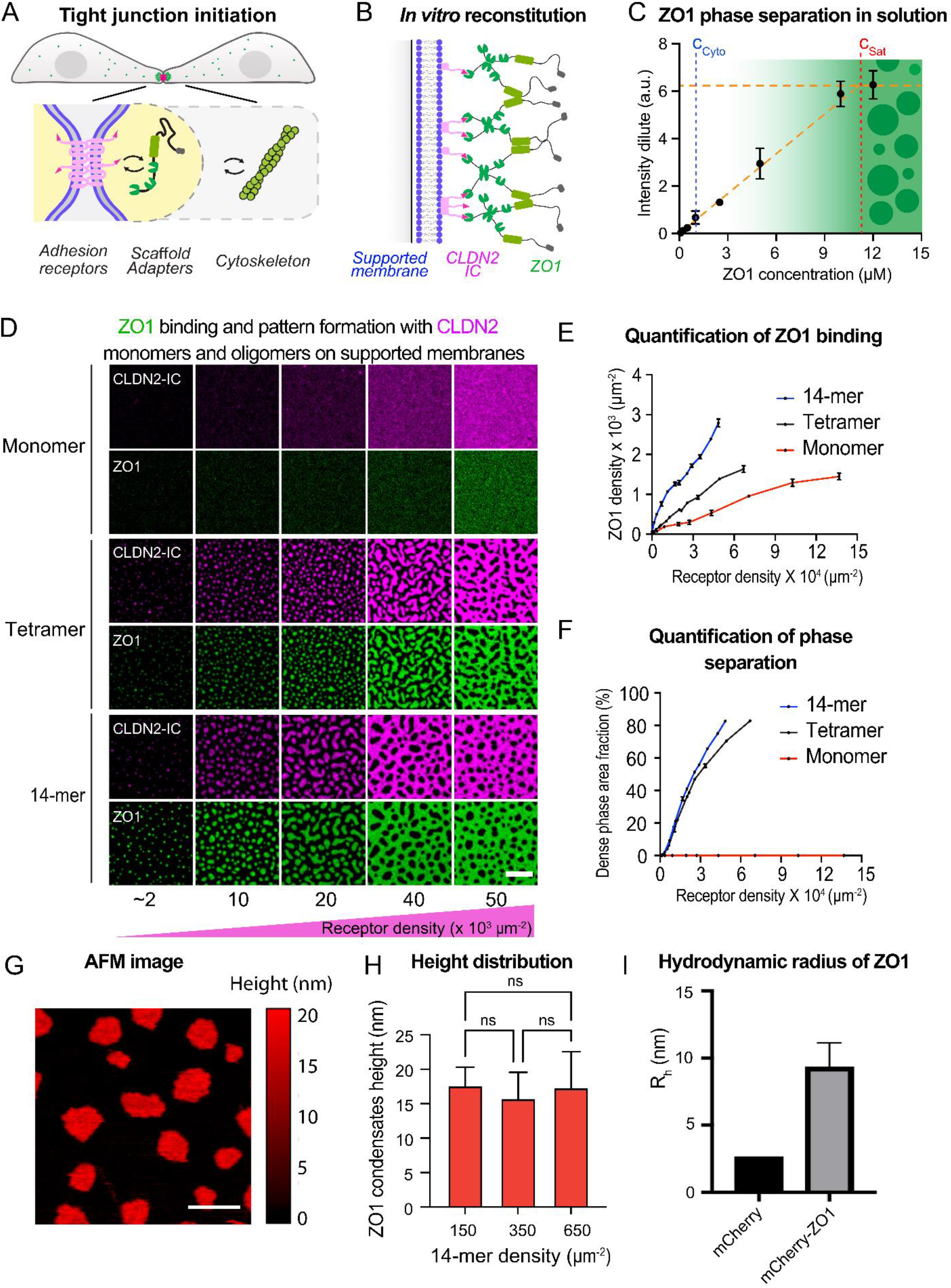
Receptor oligomerization drives ZO1 surface binding and condensation under physiological concentration. (A) Schematic of tight junction initiation with cell-cell contact in epithelial cells. (B) Schematic of the reconstitution system with supported lipid bilayers (SLBs) and tight junction components. (C) 3D saturation concentration of ZO1 (C_sat_) quantified by fluorescence intensity in dilute phase. Values shown are the mean ± SD from three independent experiments. The 3D saturation concentration (C_sat_) and cytosolic concentration (C_cyto_) are annotated. (D) Images of ZO1 surface condensates with increased receptor density of 14-mer, tetramer and monomer. 100 nM ZO1 was added to membrane-bound receptors for 15 min to form ZO1 surface condensates. Scale bar, 5μm. (E) Quantification of ZO1 membrane binding from the images in (D). Values shown are the mean ± SD from three different views. (F) Quantification of dense phase area fraction from the images in (D. Values shown are the mean ± SD from three different views. (G) AFM images of ZO1 surface condensates. ZO1 membrane condensates were formed by adding 25 nM ZO1 to 650 molecules/ μm^2^ membrane-bound 14-mer receptors for 15 min. Scale bar, 1μm. (H) Quantification of surface height from AFM images with different 14-mer densities. ZO1 membrane condensates were formed by adding 100 nM ZO1 to different amounts of membrane-bound 14-mer receptors for 15 min. Values shown are the mean ± SD from three different views. (I) Hydrodynamic radius (R_h_) of mCherry and mCherry-ZO1 proteins from FCS measurement. Values shown are the mean ± SD from 3 independent measurements.

### Reconstitution of ZO1 membrane binding and surface condensation

In our vitro reconstitution setup, ZO1 condensates formed at concentrations larger than c_sat_ = 11 μM in solution (Fig. 1C), which is close to the cytoplasmic saturation concentration measured in epithelial cells (*4*). The physiological concentration of ZO1 in the cytoplasm c_cyto_ = 690 nM is more than an order of magnitude lower than this bulk saturation concentration (*4*). Thus, at physiological ZO1 concentration, condensates cannot spontaneously form in solution. However, we found that at these subsaturated conditions ZO1 proteins can bind to membranes containing monomeric, tetrameric and 14-meric CLDN2 receptors and thereby induce phase separation at the membrane surface (Fig.1, D-F). In absence of ZO1, monomeric and oligomeric receptors were homogeneously distributed and diffused quickly (Fig. S1, C to E). While ZO1 binding to monomeric receptors was homogeneous, binding of ZO1 to tetrameric and 14-meric receptors induced formation of phase separated condensates on the membrane surface (Fig. 1D). These surface condensates were not only strongly enriched in ZO1 but also dense in receptors compared to the coexisting dilute membrane phase. Imaging the dynamics of surface condensates showed initial coarsening and fusion of condensates on a timescale of 10 min indicating liquid-like properties (Movie S1). However, over time the dynamics slowed down and condensates size typically saturated in the micrometer range (Fig. S2A). FRAP analysis showed partial exchange of ZO1 from the bulk but little diffusion in 2D (Fig. S2B), which is consistent with measurements at junctions in live cells (*4*).

To determine if the ZO1 surface condensates consist of a 2D protein monolayer or further extend into the bulk, we quantified the height of the ZO1 condensates on top of the membrane using atomic force microscopy (AFM) (Fig. 1G). AFM imaging revealed an average condensate height of 16 nm with respect to the membrane, which was largely independent of receptor surface density indicating that the structure of the condensed phase is stable over a large concentration range (Fig. 1H). Diffusion measurements of ZO1 in solution using fluorescence correlation spectroscopy showed a hydrodynamic diameter of 18 nm for a ZO1 dimer (Fig. 1I). Together, these measurements are consistent with ZO1 undergoing a surface phase transition into a condensed monolayer induced by binding to oligomerized adhesion receptors at subsaturated bulk concentrations.

### Receptor oligomerization induces robust surface condensation far below bulk saturation

To understand how ZO1 surface binding and receptor oligomerization control surface phase separation, we developed a non-equilibrium thermodynamic model (Fig. 2A). In this model, scaffold proteins in the bulk solvent can bind to receptors in the membrane composed of lipids. Accounting explicitly for the receptors bound to scaffold proteins, the minimal model corresponds to a ternary mixture in the membrane and a binary model in the bulk. We describe the interactions among all components in bulk, membrane and between bulk and membrane using a Flory-Huggins free energy; detail see SI Theory. The model can be reduced to two unknown parameters, the dilute receptor binding affinity Ω and the interaction parameter χ characterizing the interactions among ZO1 proteins that are bound to receptors. Both parameters were obtained by fitting the model at equilibrium to experimental measurements of membrane bound ZO1 proteins as a function of receptor concentration and bulk ZO1 concentration (Fig. 2, B and D).

**Fig. 2.**
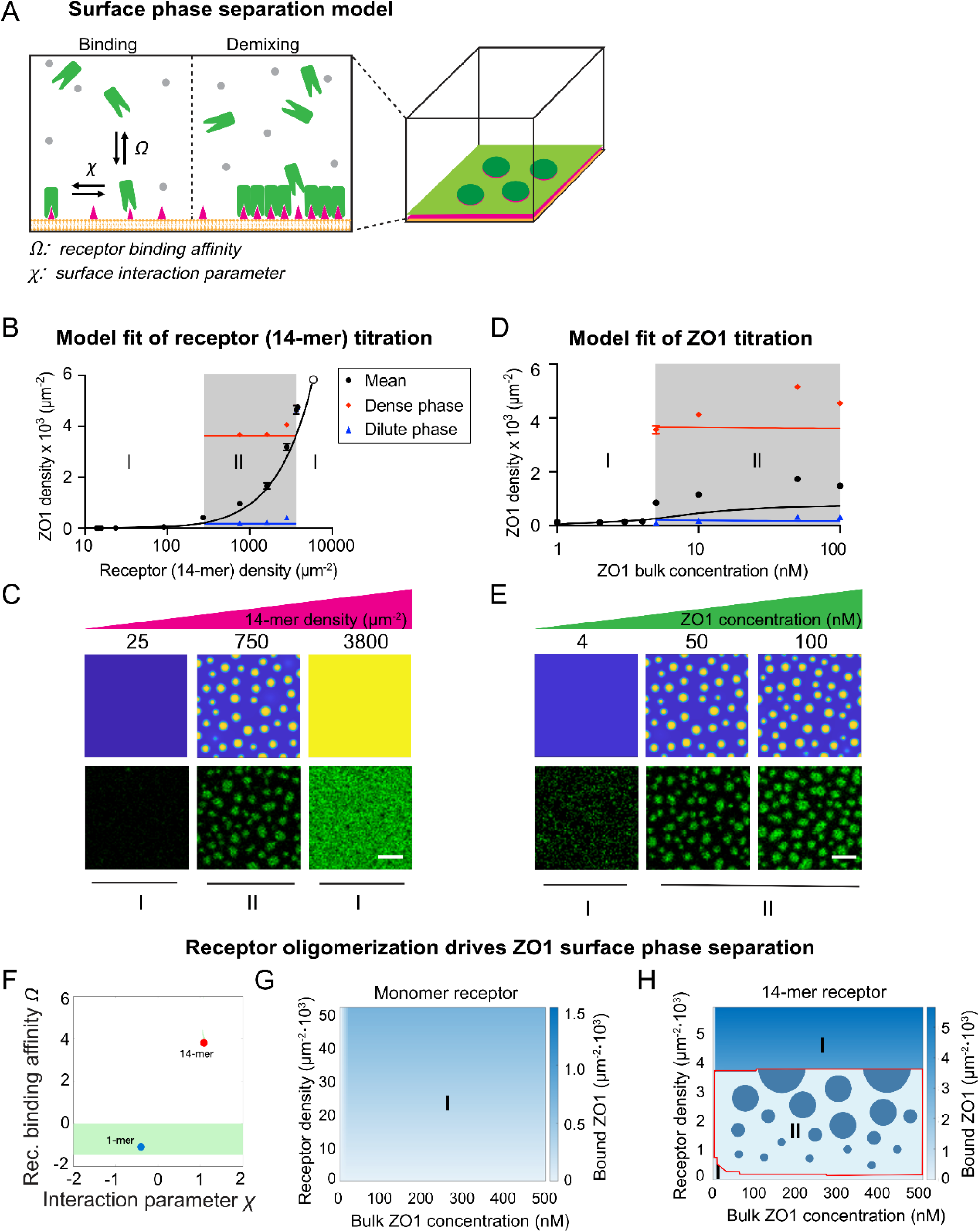
ZO1 membrane condensates form via surface phase separation under the sub-saturation regime. (A) Schematic of the surface phase separation model with specific receptor binding. Left shows the two free parameters the receptor binding affinity Ω, which controls how much ZO1 can accumulate at the surface. And the surface interaction parameter χ, which contributes to the condensation of ZO1-receptor complexes. (B) Model fitting of ZO1 density on whole membrane, in dense phase and in dilute phase when titrating the 14-mer receptor densities. Values shown are the mean ± SD from three different views. (C) Up: Snapshots of ZO1 surface condensates from simulation when titrating the 14-mer receptor density. Down: Images of ZO1 surface condensates from experiment when titrating the 14-mer receptor density. ZO1 surface condensates were formed by adding 50 nM ZO1 to the membrane functionalized with different amounts of 14-mer receptors. Scale bar, 2 μm. (D) Model fitting of ZO1 density on whole membrane, in dense phase and in dilute phase when titrating the ZO1 bulk concentration. ZO1 surface condensates were formed by adding different amounts of ZO1 to the membrane functionalized with 750 molecules/μm^2^ 14-mer receptors. Scale bar, 2 μm. (E) Up: Snapshots of ZO1 surface condensates from simulation when titrating the ZO1 bulk concentration. Down: Images of ZO1 surface condensates from experiment when titrating ZO1 bulk concentration. ZO1 surface condensates were formed by adding different amounts of ZO1 to the membrane functionalized with 750 molecules/μm^2^ 14-mer receptors. Scale bar, 2 μm. (F) Binding affinity (Ω) and interaction parameter (𝒳) with monomer and 14-mer from the fitting. In the phase diagram, the area demarcated in light green represents all parameter sets that yield Mean Square Error (MSE) values less than or equal to 130% of the minimal error observed. This zone therefore indicates a level of performance close to optimal, within an acceptable threshold of variability. (G) Phase diagram of ZO1 membrane binding with monomer receptor on membrane from simulation. Phase I shows the regime with mixed surface and mixed bulk. (H) Phase diagram of ZO1 membrane binding with 14-mer receptor on membrane from simulation. Different regimes are annotated. Phase I shows the regime with mixed surface and mixed bulk. Phase II shows the regime with demixed surface and mixed bulk.

Despite the minimal nature of the model, we found very good agreement between the model and the experimental measurements. In particular, the model recapitulated the experimental ZO1 and receptor (14-mer) concentrations at which surface condensates appear as well as the concentrations in the condensed and dilute phases (Fig. 2, B to E). Consistent with the experimental data, the model predicted a jump of the difference in ZO1 concentrations between condensates and their surrounding dilute phase. To compare the dynamics of surface condensation and growth, we chose the kinetic parameters of the model in accordance with the FRAP studies of surface condensates (Fig. S2B). In agreement with experiments, we found that above the critical concentrations surface condensates form quickly but then transition to very slow coarsening kinetics (Fig.S2A, movie S1). This slow-down is caused by depletion of receptors in the dilute phase due to co-condensation of receptors and scaffold proteins, which effectively decreases ripening fluxes among surface condensates (Fig. 2, C and E). Taken together, the model suggests that ZO1 binding to oligomeric receptors triggers a discontinuous surface phase transition at the membrane.

After successfully calibrating the model, we analyzed how the oligomerization state of the receptor affects the surface phase transition. Fitting our model to the experimental data of both receptor states (Fig. 2, B and D and Fig. S2, C, and D), we determined the change in receptor binding affinity Ω and self-interaction parameter χ of receptor bound to ZO1 for monomeric and 14-meric receptors (definitions, see SI Theory Section G). The key finding is that the binding affinity of the 14-meric receptor to ZO1 is much stronger compared to the monomeric receptor (Fig. 2F). We found that the lower binding affinity of the monomer is dominantly caused by a reduced binding fraction, i.e., only a small fraction of monomer receptors can bind to ZO1; a mechanism previously reported in Ref. (*29*). Moreover, from the fits, we obtained that the bound ZO1-14-mer complexes strongly attract each other, i.e., they have a positive self-interaction parameter (Fig. 2F). While the fitted self-interaction of the ZO1-monomer complex was lower compared to the 14mer, a wide range of self-interaction parameters fitted the experimental data equally well (green shade), indicating that in the low affinity binding regime of the monomer receptor self-interactions at the surface play a minor role (see SI Theory Fig.3(a) and its related discussion). In summary, our results suggest that receptor oligomerization favors surface phase separation via two synergistic mechanisms. Increasing the binding affinity leads to a higher surface concentration of ZO1 at constant bulk concentration. While at the same time the self-interaction of the ZO1-receptor complex lowers the saturation concentration required for membrane phase separation. Using the fitted parameters, we predicted the phase diagram for the monomeric and 14-meric receptors, which showed that there is no surface phase transition for monomeric receptors even for higher ZO1 concentrations, while there is a large region of surface phase separation for the 14-meric receptors (Fig. 2, G and H and Fig. S2, E and F). Based on the agreements between experiments and our thermodynamic model, we conclude that a surface phase transition underlies the formation of ZO1-rich surface condensation on membrane surfaces. The surface phase transition is controlled by the receptor’s oligomerization state, suggesting cell adhesion-induced oligomerization could act as the key switch for ZO1 surface condensation at cell-cell contacts.

### Selective sorting of tight junction components by ZO1 surface condensates

To study the relevance of surface condensates for tight junction assembly, we further increased the molecular complexity of our *in vitro* reconstitution (Fig. 3A). First, we asked whether ZO1 surface condensates can partition membrane receptors belonging to the adherens junction (CDH1), the tight junction (JAM-A) and the apical membrane (CRB3) (Fig. 3B). Before addition of ZO1, the intracellular domains of all receptors were well mixed on supported membranes (Fig. 3C). As before, addition of ZO1 initiated condensation of dense CLDN2-ZO1 domains. We found that the IC-domain of the tight junction receptor JAM-A was 2-fold enriched in the dense ZO1 phase. This is in line with JAM-A binding to ZO1 via PDZ2, which is independent of CLDN binding via PDZ1 (*30*) and suggests that ZO1 can simultaneously bind and crosslink different membrane receptors via its array of PDZ domains (*31*). The adherens junction receptor CDH1 was evenly distributed between the dense and dilute phase indicating that there are no direct interactions with ZO1. Surprisingly, we found that the ICD of the apical polarity receptor CRB3 was excluded from the ZO1 condensates, suggesting that segregation of apical and junctional proteins can be facilitated by condensation of ZO1 (Fig. 3, C and D). Thus, surface condensates could be an auxiliary mechanism in addition to apical polarization ensuring that cell-cell contacts are cleared from apical proteins to facilitate junction formation (*32-34*).

**Fig. 3.**
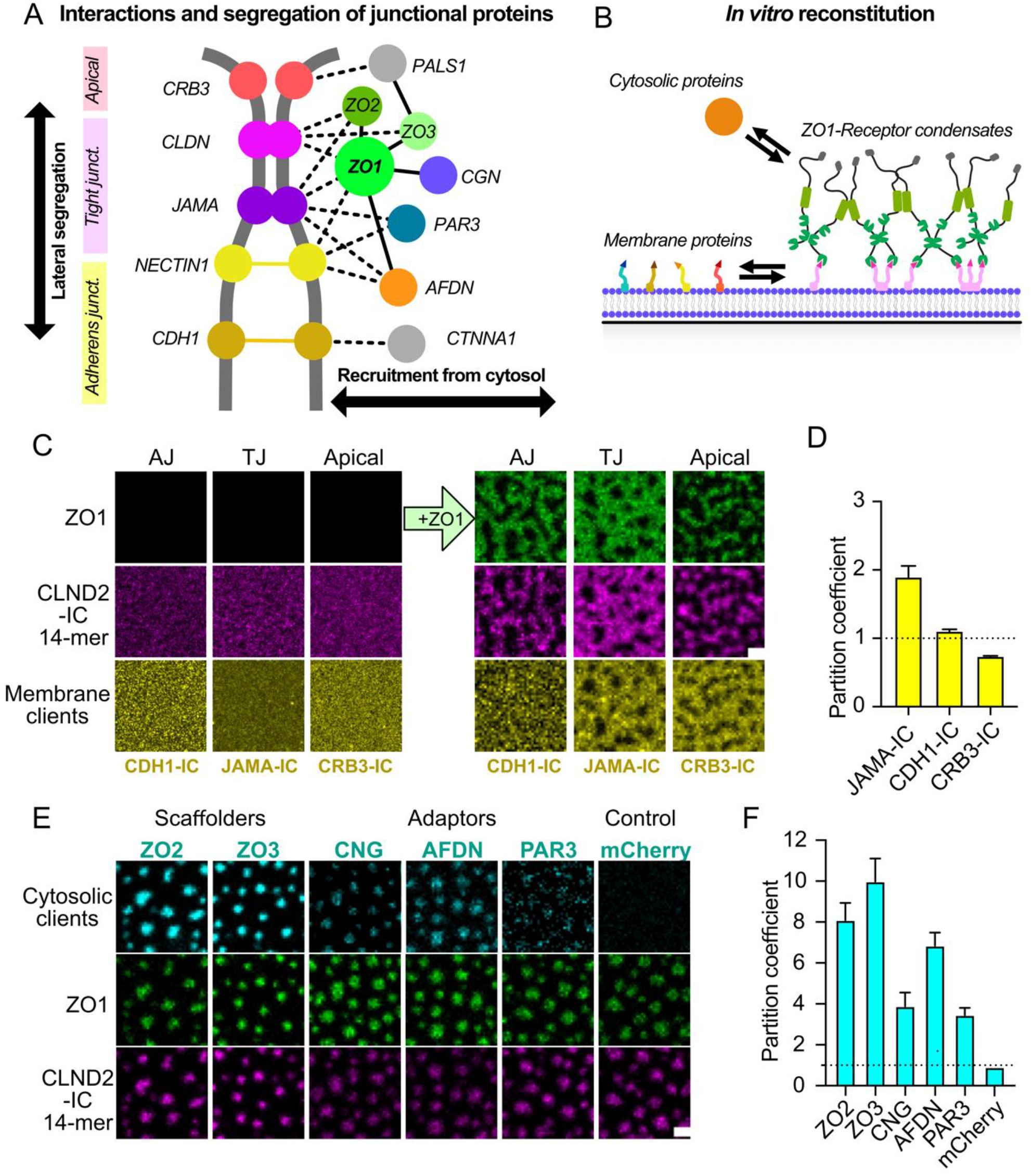
ZO1 surface condensates selectively enrich tight junction components. (A) Schematic of tight junction structures with indication of different tight junction components and non-tight junction components in lateral and apical. (B) Schematic of partition assay with different cytosolic and membrane components. (C) Left: Images of membrane functionalized with 14-mer together with JAM-A-IC, 14-mer together with E-Cad-IC, and 14-mer together with CRB3-IC. Right: Images of different receptors partitioned into or segregated with ZO1 surface condensates. ZO1 surface condensates were formed by adding 100 nM ZO1 for 15 min. Scale bar, 2 μm. (D) Partition coefficient quantification of different receptors into ZO1 surface condensates from images in (C). Values shown are the mean ± SD from 3 different views. (E) Images of cytosolic tight junction components partitioning into ZO1 surface condensates. mCherry protein was used as a negative control. ZO1 membrane condensates were formed by adding 100 nM ZO1 to 680 molecules/μm^2^ membrane-bound 14-mer for 15 min. Images were taken after adding 100 nM client proteins to ZO1 surface condensates for 15 min. Scale bar, 2 μm. (F) Partition coefficient quantification of cytosolic tight junction components into ZO1 surface condensates from images in (E). Values shown are the mean ± SD from 3 different views.

In addition to membrane receptors, we investigated the recruitment of cytosolic scaffold and adapter proteins (Fig. 3B). As predicted by our previous work studying ZO1 condensation in solution (*4*), we found that the two ZO1 homologues ZO2 and ZO3 strongly partitioned in ZO1 surface condensates. The ZO1 binding proteins CGN, AFDN and PAR3 also became significantly enriched in the condensed phase, while the control protein mCherry showed neutral partitioning (Fig. 3, E and F). Together, these results established that ZO1 surface condensation drives sorting of key junctional components such as adhesion receptors and adapter proteins required for tight junction assembly into concentrated membrane domains which exclude the apical polarity proteins. While the colocalization of junctional proteins and the segregation of apical proteins *in vitro*, recapitulated to some degree the molecular organization of mature junctions in cells (*1*), the overall distribution and shape of the surface condensates was not changed and remained in a state that matched isolated condensates during the initial phase of junction assembly (*4*).

### Local actin polymerization drives surface condensates elongation

To understand how nascent ZO1 surface condensates are reorganized into a belt structure resembling the structure of tight junctions, we investigated the interplay of the condensates with the actin cytoskeleton (Fig. 4A). Mature tight junctions are surrounded by a sub-apical belt of filamentous actin and perturbations of the actin cortex have been shown to disrupt junction structure and induce opening of the *trans-*epithelial barrier (*35, 36*). The junctional scaffold proteins ZO1, ZO2, CGN and AFDN contain actin-binding-domains (ABD), which have been suggested to provide the connection between the membrane receptors and the actin cortex (*31, 35*). However, how actin binding is mechanistically related to junction formation is poorly understood. We first measured actin polymerization in the presence of 200 nM ZO1 in solution using G-actin labeled with pyrene (Fig. 4B). We found no significant influence of ZO1 on actin polymerization in solution, while a positive control (WASP+Arp2/3) showed strong increase. Next, we added fluorescently labeled G-actin to ZO1-CLDN2 membrane condensates. Under these conditions, actin was sequestered to surface condensates and rapidly polymerized into a membrane bound network (Fig.4, A and C, and movie S2). While no actin polymerization was observed on membranes without surface bound ZO1 or receptors (Fig. S3, C and D). Strikingly, actin polymerization induced reshaping of isolated condensates into a connected strand network with ZO1 and CLDN2 colocalizing with actin fibers. During the reshaping process ZO1 and CLDN2 surface concentrations remained constant while actin concentrations increased monotonically (Fig 4C). The initial elongation rate of the actin-condensates was 0.5 ± 0.1 μm/min (Fig. 4D), which is close to previously determined polymerization rates of single actin fibers (*37*). Over time, actin filaments formed bundles by polymerization of new filaments along existing fibers (Fig. 4, E and F, and movie S3). Repeating the same experiment with a ZO1 mutant lacking the actin-binding-domain failed to induce actin polymerization (Fig. 4, G and H), despite similar surface phase separation properties of the ΔABR mutant as WT-ZO1 (Fig. S3, A and B). Quantification showed that actin became 5-fold enriched into WT-condensates while the ZO1-ΔABR was still able to enrich actin it was 2-fold reduced (Fig. 4I). In addition, comparing actin recruitment to non-phase separated ZO1 bound to monomeric receptors on the membrane, showed that both actin polymerization and bundling were significantly reduced (Fig. S3, E and F).

**Fig. 4.**
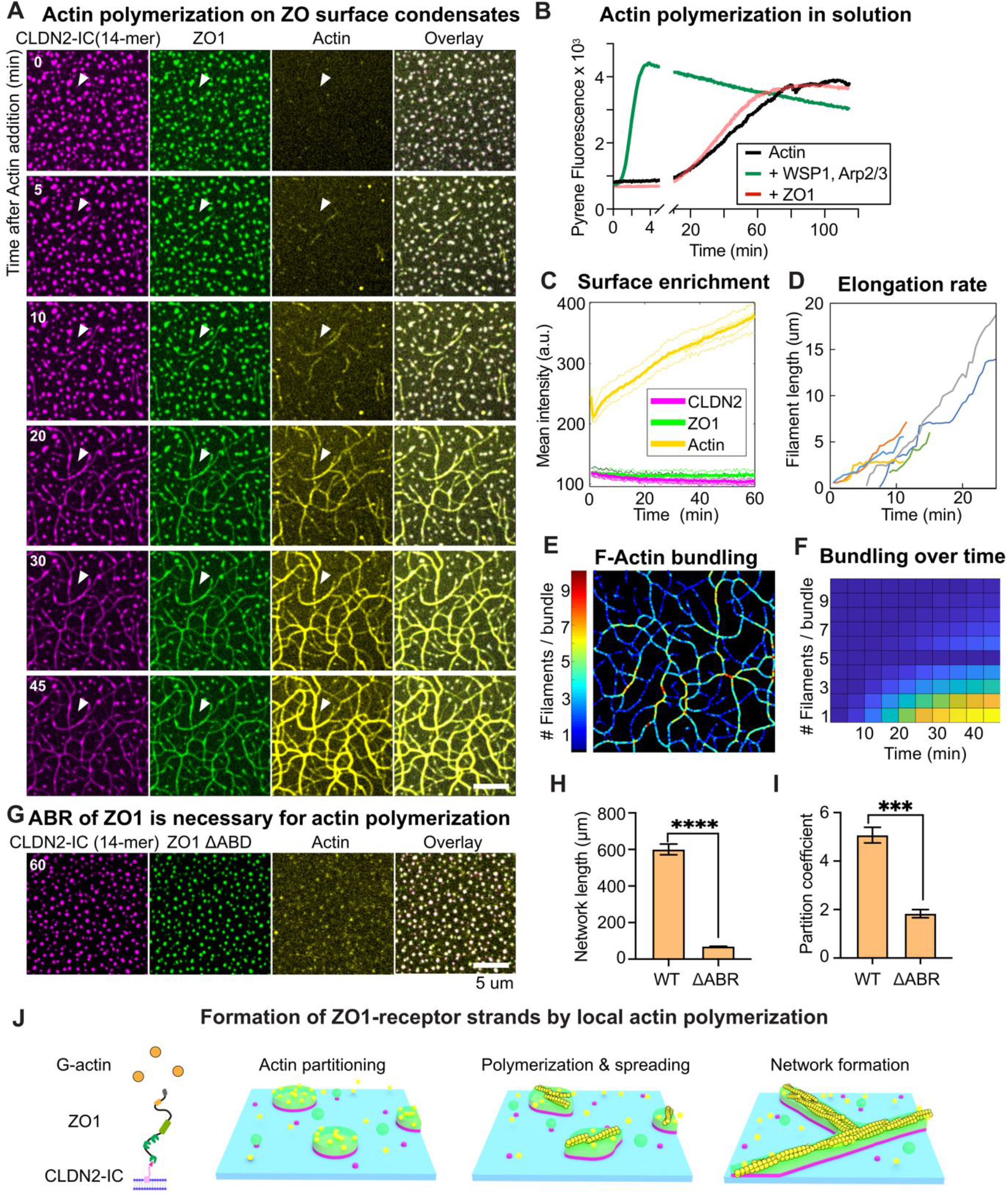
Local actin polymerization drives elongation of junctional condensates. (A) Time series of local actin polymerization and ZO1 surface condensates deformation. ZO1 surface condensates were formed by adding 200 nM ZO1 to 450 molecules/μm^2^ membrane-bound 14-mer-DyLight 550 for 10min. Image series was taken after adding 3 uM G-actin-Alexa Fluor 647 to ZO1 membrane condensates every 30 s for 60 min. The white arrowhead indicates one example of actin filament polymerization and ZO1 surface condensates spreading. Scale bar, 2 μm. (B) Pyrene actin polymerization assay in tube with 200 nM ZO1 or 100 nM Arp2/3 and 100 nM WASP1. (C) Quantification of ZO1, 14-mer receptor and actin fluorescence on membrane during time from the time series in (A). (D) Quantification of actin filaments growth during time from the time series in (A). (E) Color map of F-actin bundling degree after adding 3 uM G-actin-Alexa Fluor 647 for 60 min. (F) F-actin bundling degree during time from the time series in (A). (G) Actin local polymerization assay with ZO1-ΔABR surface condensates. ZO1-ΔABR surface condensates were formed by adding 200 nM ZO1-ΔABR protein to the membrane functionalized with 450 molecules/μm^2^ 14-mer for 10 min. Images were taken after adding 3 uM G-actin-Alexa Fluor 647 for 60 min. Scale bar, 10 μm. (H) Quantification of actin network length with ZO1 surface condensates formed with ZO1-WT and ZO1-ΔABR. Values shown are the mean ± SD from 3 different views. Unpaired t-test was performed to determine the significance of the difference. P = 0.0001. (I) Partition coefficient of actin signal into the ZO1 surface condensates formed with ZO1-WT and ZO1-ΔABR. Values shown are the mean ± SD from 3 different views. Unpaired t-test was performed to determine the significance of the difference. P < 0.0001. (J) Schematic of local actin polymerization, ZO1 surface condensates spreading and tight junction like network formation on membrane.

Together, the results show that surface condensation of ZO1-CLDN2 is sufficient to sequester actin from solution and induce polymerization in an ZO1-ABR dependent manner (Fig. 4J). This suggests that the combination of weak actin binding affinity of ZO1 (*36*) together with the high density of actin binding sites in the surface condensates, provides robust spatiotemporal control for the assembly of an actin cortex at nascent cell-cell adhesion sites. In addition, the liquid-like material properties of the condensed phase enable its spreading along the growing actin filaments, which results in the formation of continuous CLDN2-ZO1-Actin strands resembling the molecular organization of the tight junction belt. Interestingly, during the formation of the reconstituted junctional strands actin filaments grow as bundles (Fig. 4, E and F). Actin bundling, in contrast to branching, leads to elongation of condensates along a line and connects the condensates into a sparse network with both its connectivity and complexity are constrained by the initial condensate density (Fig. S4). Thus, bundling could be an important feature of ZO1 condensates that promotes formation of a narrow subapical tight junction belt and prevents ectopic expansion of the tight junction into the lateral membrane domain.

Finally, we compared our *in vitro* reconstitution results to actin dependent tight junction assembly in epithelial tissue culture. To this end we prepared an MDCKII cell line that allowed us to follow the dynamics of endogenous ZO1 (NeonGreen tagged), and F-actin (Utrophin labeled) during junction assembly after perturbations via actin depolymerization by Latrunculin-A (LatA) or calcium depletion (*4, 38*). In confluent MDCK monolayers ZO1 and F-actin colocalized at the TJ-belt (Fig. 5A). Addition of 1 μM LatA let to rapid breakage of the TJ-belt and induced formation of condensates at the cell-cell interface resembling the initial ZO1 surface condensates in our *in vitro* experiments. Wash out of LatA induced actin polymerization and reformation of the junctional belt (Fig. 5B). Similar to our *in vitro* experiments, actin extended together with ZO1 condensates and connected single condensates into a continuous belt. Repeating these experiments using a Calcium switch perturbation to reassemble junctions, resulted in similar behavior (Fig. 5B). Comparison of the elongation rates of junctional condensates quantified in MDCK tissue after LatA and Calcium switch with the elongation rates of our *in vitro* reconstitution assay showed a striking agreement (Fig. 5C). These results suggests that actin polymerization is the rate limiting process for tight junction belt elongation.

**Fig. 5.**
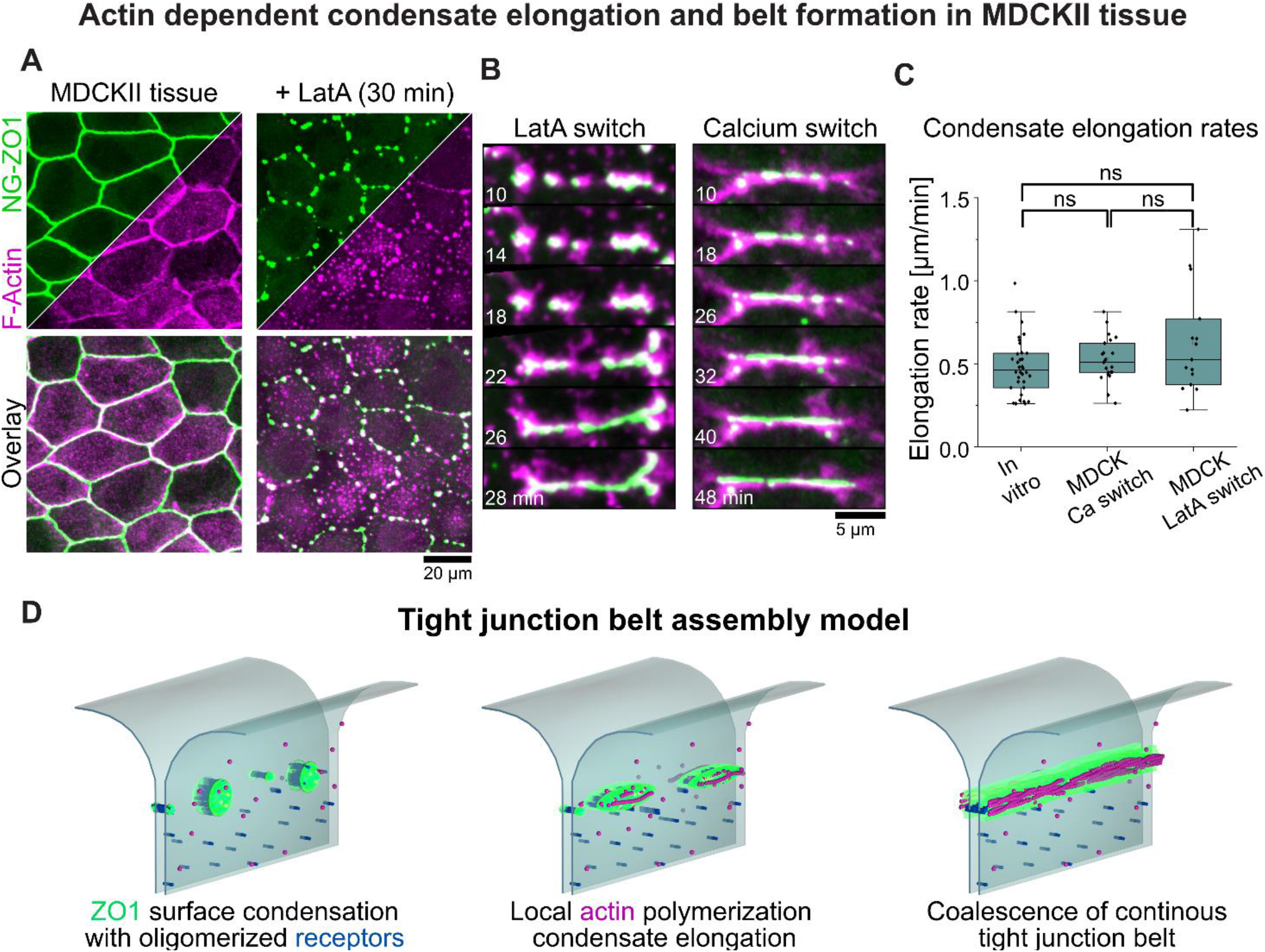
Actin dependent elongation of junctional condensates in epithelial tissue. (A) Colocalization of ZO1 and F-actin in confluent monolayers of MDCK cells. Endogenous ZO1 was visualized via fusion NeoGreen (NG). F-actin was visualized via the binding protein UtrCH fused to a SNAP-tag and stained with SiR-SNAP. Addition of 1μM Latrunculin-A (LatA) induces actin depolymerization and results in the formation of junction condensates enriched in ZO1 and actin. (B) Junction assembly assay using the same cell line described in (A). Left panel: LatA was wash out and ZO1 and F-actin were imaged during the reformation of junctional belts. ZO1 and F-actin colocalize during elongation of the junctional condensates. Right panel: Reformation of junction after calcium switch showed similar dynamics as the LatA switch. (C) Quantification of the elongation speed of individual condensates from the in vitro experiments shown in figure 4 and the LatA and Ca switch experiments. Elongation rates have a median of 0.5 ± 0.2 μm/min for in vitro experiments, 0.5 ± 0.1 μm/min for calcium switch and 0.6 ± 0.3 μm/min for the LatA switch. (D) Model of tight junction belt assembly based on the experimental and theoretical results. ZO1 is recruited to cell-cell adhesion sites, by binding to oligomeric receptors. Multivalent interaction of ZO1 drives surface condensation which induces partitioning of receptors and cortical proteins including actin. ZO1 condensates facilitate local actin-polymerization, which drives elongation of condensates and results in coalescence into continuous junctional belts.

## Conclusion

To summarize, we reconstituted the self-assembly of the intracellular part of the tight junction belt in an *in vitro* system, which enabled control of the concentrations of the components, quantification of molecular interactions, and observation of emergent behavior in ways that are difficult to achieve using intact tissue. Combining the experimental results with thermodynamic modeling, we found that a surface phase transition is triggered when cytoplasmic ZO1 binds to oligomeric adhesion receptors in the membrane. Exploring the coupling between bulk and surface interactions, revealed that surface condensation is largely independent of the cytoplasmic concentration of ZO1 and is rather controlled by receptor concentration and oligomerization. Thus, the interactions are tuned such that condensation in the bulk is suppressed due to sub-saturation but condensation at the membrane can be locally triggered by receptor enrichment and oligomerization, which are both occurring at cell-cell contact sites (*39, 40*). We suggest that this mechanism provides robust spatiotemporal control to couple extracellular adhesion signals to intracellular assembly of the tight junction scaffold and possibly other adhesion junctions (*16, 41*).

Akin to other membrane condensates (*14-16*), we found that ZO1 condensation can create an environment that facilitates local biochemical reactions. ZO1 condensates enriched key junctional components, excluded apical proteins and facilitate actin polymerization and filament bundling. ZO1 condensation mediated actin polymerization drove a morphological transition from isolated domains into a continuous network of receptor-ZO1-actin strands. This reconstituted transition was remarkably similar in terms of morphology and kinetics to the junction assembly process in cells. While to create the junctional belt in cells additional mechanism will be required to align the elongating junctional condensates along the apical interface (*42*), our results suggest that a key driver for the elongation itself is the interplay between local actin polymerization and the cohesive forces of the ZO1 condensate. In contrast to other membrane condensates that interact with or nucleate actin filaments (*14, 15*), low affinity actin binding is directly inbuilt into the main scaffolder ZO1, which provides direct coupling between the surface phase transition and actin nucleation without partitioning of other actin nucleators. However, there are many additional actin binding proteins recruited to the junctional condensates, which may add robustness to system (*43*) and/or provide additional functions such as mechano-sensing (*44, 45*). In addition to the actin dependent mechanism uncovered in this work, it will be important to understand how ZO1 condensates relate to the polymerization of Claudin receptors, which form the actual diffusion barrier.

More generally, our work shows how cells control and exploit the collective properties of protein condensates to actively assemble and shape complex biological structures. Recent examples have shown that interactions of protein condensates with biological surfaces such as DNA, cytoskeletal filaments or lipid membranes can drive mesoscale shape changes such as DNA compaction, cortex formation and membrane folding and budding (*24, 46, 47*). Here we uncovered how cells convert punctual extracellular adhesion cues into continuous tight junction belts by exploiting surface phase transitions coupled to actin polymerization. We envision that our reconstitution platform in combination with our thermodynamic model will provide a template to uncover the assembly mechanism of other adhesion complexes and surface coupled cellular structures in general.

## Acknowledgments

D.S. acknowledges support from PEPC facility at MPI-CBG for protein purification. D.S. acknowledges support from Dr. Jens Ehrig of the Molecular Imaging and Manipulation Facility, a core facility of the CMCB at TU Dresden for AFM measurement. C.W. and X.Z. acknowledge insightful discussions about physical modeling with Frank Jülicher. We are thankful for the discussions and helpful comments on the manuscript by Christoph Zechner.

## Author contributions

A.H. and D.S. conceived the experimental part of the project. C.W. and X.Z. conceived the physical model. D.S. purified proteins and performed all experiments. X.Z. implemented and fitted the model. T.W. performed the actin polymerization assay in solution and helped with actin polymerization on SLBs. C.M. established stable MDCKII cell lines. D.S., A.H., X.Z. analyzed the experimental data. A.H., C.W., D.S., X.Z and A.A.H. wrote the manuscript.

## Data and materials availability

All data are available in the main text, the supplementary materials or supplementary information on theoretical model.

## Supplementary Materials

Materials and Methods

Figs. S1 to S4

Table S1

References (48–49)

Movies S1 to S3

## Supplementary Information on Theoretical Model

A. Model for protein binding to receptors in a membrane

B. Thermodynamics and molecular interactions

C. Equilibrium thermodynamics

D. Binding fraction

E. Non-equilibrium thermodynamics with membrane binding

F. Parameter choices and parameters obtained from fitting to the experimental data

G. Derivation of dilute binding affinity given in Eq. (23)

H. Kinetics of membrane phase separation

References

## References

1. C. Zihni, C. Mills, K. Matter, M. S. Balda, Tight junctions: from simple barriers to multifunctional molecular gates. Nat Rev Mol Cell Biol 17, 564–580 (2016).

2. J. M. Anderson, C. M. Van Itallie, Physiology and function of the tight junction. Cold Spring Harb Perspect Biol 1, a002584 (2009).

3. S. Citi, Intestinal barriers protect against disease. Science 359, 1097–1098 (2018).

4. O. Beutel, R. Maraspini, K. Pombo-Garcia, C. Martin-Lemaitre, A. Honigmann, Phase Separation of Zonula Occludens Proteins Drives Formation of Tight Junctions. Cell 179, 923-+ (2019).

5. K. Umeda et al., ZO-1 and ZO-2 independently determine where claudins are polymerized in tight-junction strand formation. Cell 126, 741–754 (2006).

6. C. Schwayer et al., Mechanosensation of Tight Junctions Depends on ZO-1 Phase Separation and Flow. Cell 179, 937–952 e918 (2019).

7. C. P. Brangwynne et al., Germline P Granules Are Liquid Droplets That Localize by Controlled Dissolution/Condensation. Science 324, 1729–1732 (2009).

8. A. Patel et al., A Liquid-to-Solid Phase Transition of the ALS Protein FUS Accelerated by Disease Mutation. Cell 162, 1066–1077 (2015).

9. D. Sun, R. Wu, J. Zheng, P. Li, L. Yu, Polyubiquitin chain-induced p62 phase separation drives autophagic cargo segregation. Cell Res 28, 405–415 (2018).

10. C. P. Brangwynne, T. J. Mitchison, A. A. Hyman, Active liquid-like behavior of nucleoli determines their size and shape in Xenopus laevis oocytes. P Natl Acad Sci USA 108, 4334–4339 (2011).

11. A. Boija et al., Transcription Factors Activate Genes through the Phase-Separation Capacity of Their Activation Domains. Cell 175, 1842-+ (2018).

12. A. W. Fritsch et al., Local thermodynamics govern formation and dissolution of Caenorhabditis elegans P granule condensates. Proc Natl Acad Sci U S A 118, (2021).

13. A. A. Hyman, C. A. Weber, F. Julicher, Liquid-liquid phase separation in biology. Annu Rev Cell Dev Biol 30, 39–58 (2014).

14. X. Su et al., Phase separation of signaling molecules promotes T cell receptor signal transduction. Science 352, 595–599 (2016).

15. M. Zeng et al., Reconstituted Postsynaptic Density as a Molecular Platform for Understanding Synapse Formation and Plasticity. Cell 174, 1172–1187 e1116 (2018).

16. Y. Wang et al., LIMD1 phase separation contributes to cellular mechanics and durotaxis by regulating focal adhesion dynamics in response to force. Dev Cell 56, 1313–1325 e1317 (2021).

17. P. G. Degennes, Wetting - Statics and Dynamics. Rev Mod Phys 57, 827–863 (1985).

18. J. W. Cahn, Critical-Point Wetting. J Chem Phys 66, 3667–3672 (1977).

19. J. E. Rutledge, P. Taborek, Prewetting Phase-Diagram of He-4 on Cesium. Phys Rev Lett 69, 937–940 (1992).

20. S. Chandavarkar, R. M. Geertman, W. H. Dejeu, Observation of a Prewetting Transition during Surface Melting of Caprolactam. Phys Rev Lett 69, 2384–2387 (1992).

21. H. Kellay, D. Bonn, J. Meunier, Prewetting in a Binary-Liquid Mixture. Phys Rev Lett 71, 2607–2610 (1993).

22. X. P. Zhao, G. Bartolucci, A. Honigmann, F. Julicher, C. A. Weber, Thermodynamics of wetting, prewetting and surface phase transitions with surface binding. New J Phys 23, (2021).

23. J. A. Morin et al., Sequence-dependent surface condensation of a pioneer transcription factor on DNA. Nature Physics 18, 271-+ (2022).

24. T. Quail et al., Force generation by protein-DNA co-condensation. Nature Physics 17, 1007-+ (2021).

25. H. Sasaki et al., Dynamic behavior of paired claudin strands within apposing plasma membranes. P Natl Acad Sci USA 100, 3971–3976 (2003).

26. J. Zhao et al., Multiple claudin-claudin cis interfaces are required for tight junction strand formation and inherent flexibility. Communications Biology 1, (2018).

27. J. Liu et al., Conformational specificity of the Lac repressor coiled-coil tetramerization domain. Biochemistry-Us 46, 14951–14959 (2007).

28. B. M. Collins et al., Homomeric ring assemblies of eukaryotic Sm proteins have affinity for both RNA and DNA - Crystal structure of an oligomeric complex of yeast SmF. Journal of Biological Chemistry 278, 17291–17298 (2003).

29. S. Erlendsson et al., Mechanisms of PDZ domain scaffold assembly illuminated by use of supported cell membrane sheets. Elife 8, (2019).

30. K. Ebnet, C. U. Schulz, M. K. Meyer Zu Brickwedde, G. G. Pendl, D. Vestweber, Junctional adhesion molecule interacts with the PDZ domain-containing proteins AF-6 and ZO-1. J Biol Chem 275, 27979–27988 (2000).

31. F. Rouaud et al., Scaffolding proteins of vertebrate apical junctions: structure, functions and biophysics. Biochim Biophys Acta Biomembr 1862, 183399 (2020).

32. K. Shin, S. Straight, B. Margolis, PATJ regulates tight junction formation and polarity in mammalian epithelial cells. J Cell Biol 168, 705–711 (2005).

33. K. Sasaki et al., Shank2 Binds to aPKC and Controls Tight Junction Formation with Rap1 Signaling during Establishment of Epithelial Cell Polarity. Cell Reports 31, (2020).

34. Q. Wang, B. Margolis, Apical junctional complexes and cell polarity. Kidney Int 72, 1448–1458 (2007).

35. S. Citi, The mechanobiology of tight junctions. Biophys Rev 11, 783–793 (2019).

36. B. Belardi et al., A Weak Link with Actin Organizes Tight Junctions to Control Epithelial Permeability. Dev Cell 54, 792–804 e797 (2020).

37. J. R. Kuhn, T. D. Pollard, Real-time measurements of actin filament polymerization by total internal reflection fluorescence microscopy. Biophys J 88, 1387–1402 (2005).

38. M. Melak, M. Plessner, R. Grosse, Actin visualization at a glance. Journal of Cell Science 130, 525–530 (2017).

39. T. Otani et al., Claudins and JAM-A coordinately regulate tight junction formation and epithelial polarity. J Cell Biol 218, 3372–3396 (2019).

40. H. Gonschior et al., Nanoscale segregation of channel and barrier claudins enables paracellular ion flux. Nat Commun 13, 4985 (2022).

41. L. B. Case, M. De Pasquale, L. Henry, M. K. Rosen, Synergistic phase separation of two pathways promotes integrin clustering and nascent adhesion formation. Elife 11, (2022).

42. K. Pombo-García, C. Martin-Lemaitre, A. Honigmann, (2022).

43. M. A. Odenwald et al., The scaffolding protein ZO-1 coordinates actomyosin and epithelial apical specializations in vitro and in vivo. J Biol Chem 293, 17317–17335 (2018).

44. M. Cordenonsi et al., Cingulin contains globular and coiled-coil domains and interacts with ZO-1, ZO-2, ZO-3, and myosin. J Cell Biol 147, 1569–1582 (1999).

45. S. Paschoud, L. Guillemot, S. Citi, Distinct domains of paracingulin are involved in its targeting to the actin cytoskeleton and regulation of apical junction assembly. J Biol Chem 287, 13159–13169 (2012).

46. V. T. Yan, A. Narayanan, T. Wiegand, F. Julicher, S. W. Grill, A condensate dynamic instability orchestrates actomyosin cortex activation. Nature 609, 597–604 (2022).

47. F. Yuan et al., Membrane bending by protein phase separation. Proc Natl Acad Sci U S A 118, (2021).

