## Supplemental materials for "Assembly of tight junction belts by surface condensation and actin elongation"

**The PDF file includes:**

Materials and Methods
Figs. S1 to S4
Table S1
References (46-47)

**Other Supplementary Materials for this manuscript include the following:**

Movies S1 to S3
Supplementary Information on Theoretical Model

### Materials and Methods

Generation of monomeric, tetrameric, and 14-meric receptors

We used the last 7 amino acids from the human CLDN2 C-terminal as the receptor for recruiting ZO1 protein to the membrane. To make the receptor easier to purify, we fused a SNAP-tag before the 7 amino acids to get a recombinant protein SNAP-CLDN2-C7 (referred to as monomer) as a monomeric receptor. We used GCN4 and ySMF domains, which are known to form stable tetramer and 14-mer, to get recombinant proteins GCN4-CLDN2-C7 (referred to as tetramer) and ySMF-CLDN2-C7 (referred to as 14-mer) as tetrameric and 14-meric receptors.

Protein expression and purification of scaffold and adapter proteins

For ZO1, ZO2, ZO3, CGN, AFDN, and PAR3, we used insect cells to recombinantly express proteins using the baculovirus expression system. SF9-ESF *S. frugiperda* cells were infected with respective baculoviruses, and then cultured at 27 °C in ESF 921 insect cell culture medium supplemented with 2 % fetal bovine serum for 3 days. Cell pellets were then collected, washed, and resuspended with a lysis buffer (20 mM Hepes, pH 7.4, 500 mM NaCl, 1 mM $MgCl_2$, 1 x protease inhibitor cocktail, 1x benzonase). Cell pellets were broken with LM20 Microfluidizer, 5,000 psi, 2 runs, and clarified by centrifugation at 17,000 g, at 4 °C, for 30 min. Proteins were then purified with metal ion affinity chromatography (Ni-NTA) resin (IMAC, 5 mL HiTrap Chelating, GE Healthcare), and followed by amylose resin (NEB). Finally, size exclusion chromatography was performed with the Superose 6 column on AKTA pure FPLC system (GE). Proteins were collected, aliquoted, and frozen in liquid nitrogen and stored at -80 °C in 20 mM Hepes, pH 7.4, 500 mM NaCl, 1 mM DTT, and 5 % Glycerol. His and MBP tag was cleaved and removed with amylose resin before being used for respective assays.

Protein expression and purification of receptor proteins

For monomer, tetramer, 14-mer, mCherry-CDH1-IC, JAM-A-IC and mCherry-CRB3-IC, we used *Escherichia coli* Rosetta cells to recombinantly express proteins. Proteins were expressed in the *Escherichia coli* Rosetta cells in LB medium at 16 °C overnight and induced with 0.2 mM IPTG when $OD_{600}$ reached 0.6 - 0.8. All the following steps were carried out at 4 °C. Cell pellets were collected, washed and resuspended with a lysis buffer (20 mM Hepes, pH 7.4, 500 mM NaCl, 1 mM $MgCl_2$, 1 x protease inhibitor cocktail, 1x benzonase). Cell pellets were broken with LM20 Microfluidizer, 20,000 psi, 2 runs, and clarified by centrifugation at 17,000 g, at 4 °C, for 30 min. Proteins were then purified with metal ion affinity chromatography (Ni-NTA) resin (IMAC, 5mL HiTrap Chelating, GE Healthcare) and followed by size exclusion chromatography with Superdex 200 increase 10/300 GL column on AKTA pure FPLC system (GE). Proteins were collected, aliquoted, and frozen in liquid nitrogen and stored at -80°C in 20mM Hepes, pH 7.4, 500 mM NaCl, 1 mM DTT, and 5 % Glycerol.

Protein fluorescence labeling

Monomer, tetramer, 14-mer, and JAM-A-IC were labeled with DyLight550 NHS Ester (Thermo Scientific, Product No.62265) or DyLight 650 NHS Ester (Thermo Scientific, Product No.62262) dye. Highly purified proteins were prepared in 0.05 M sodium borate buffer at pH 8.5 (Thermo Scientific BupH Borate Buffer Packs, Product No. 28384). DyLight NHS Ester dyes were dissolved in DMSO. When labeling the protein, the dye and protein were mixed at a molar ratio of 1:1, and incubated at RT for 1hr. Free dye was removed from the protein by desalting columns (Thermo Scientific Zeba Spin Desalting Columns, Product No. 89882) with buffer containing 20 mM Hepes, pH 7.4, 500 mM NaCl, 1 mM DTT, 5% Glycerol. Fluorescence labeling efficiency was measured with Nanodrop 2000 (ThermoFisher). The labeled and unlabeled proteins were mixed to get a final stock of 5 % labeled.

Microscopy

FRAP, FCS, and confocal imaging were performed on a commercial confocal STED microscope (Abberior Instruments, Göttingen, Germany) with pulsed laser excitation (490 nm, 560 nm, 640 nm, 40MHz) and 60x water or 100x oil objectives (Olympus). MDCKII cells were imaged on a Nikon spinning disk microscope with a 60x water objective and a living cell imaging system supplying $CO_2$ and heating.

Fluorescence recovery after photobleaching (FRAP)

Regions of interest (ROI) were bleached using a 405 nm diode with 1.5 mW with 100 µs pixel dwell time. Pre-bleached and post-bleached images were acquired with 490 nm, 560 nm, and 640 nm laser with a frame rate of 1 s for 10 min. Recovery data were normalized to a reference ROI outside the bleached area. FRAP traces were evaluated and fitted in GraphPad Prism software.

Fluorescence correlation spectroscopy (FCS)

mGFP, 5 % DyLight 550 and 5% DyLight650 labeled 14-meric receptors were anchored to SLBs via His10/His6 tag and DGS-NTA(Ni) interaction and excited with 490 nm, 560 nm and 640 nm pulsed laser (Olympus 60x NA = 1.2 water objective). Fluorescence fluctuations were recorded with a time resolution of 500 ns for 10 s. Auto-correlation of the photon traces was performed in MATLAB using a multiple tau correlator. The resulting correlation curves were fitted according to the standard 2D diffusion model including one triplet component using MTALAB (Elson, 2011). The mean particle number N was obtained from the fitting. mGFP and ZO1-mGFP were excited with a 490 nm laser in the bulk. The resulting correlation curves were fitted according to the standard 3D diffusion model including one triplet component using MTALAB (Elson, 2011). The comparison of brightness between the two molecules was calculated by the counts per molecule.

*Calibration curve generations*

FCS was performed on SLBs functionalized with different amounts of mGFP, 14-mer-DyLight 650 and 14-mer-DyLight 550. Mean particle numbers were obtained from FCS fitting. Confocal images were taken for the SLBs functionalized with mGFP, 14-mer-DyLight 650, or 14-mer-DyLight 550 for extracting the mean fluorescence intensity. The mean fluorescence intensity of mGFP was normalized to ZO1-mGFP by the brightness comparison obtained from FCS measurement in the bulk. Calibration curves were obtained by plotting mean particle numbers as a function of fluorescence mean intensity.

*Hydrodynamic radius ($R_h$) measurement*

FCS measurements were performed on mCherry and ZO1-mCherry proteins with a 560 nm pulsed laser under the same buffer condition to get the diffusion coefficient. $R_h$ of the globular protein mCherry protein is known as ~2.6 nm. The Stokes-Einstein equation was used to get the $R_h$ of ZO1-mCherry protein by plugging the diffusion coefficient and $R_h$ of mCherry proteins into it.

<u>3D phase separation assay and saturation concentration determination</u>

MBP tag was cleaved by adding 3C proteases to ZO1-GFP-3C-MBP protein stock at RT for 3 hrs. Protein stock was then diluted to a 3D phase separation buffer (20 mM Hepes, pH 7.4, 150 mM NaCl) to a series of concentrations. Images were taken with a confocal microscope (Olympus 60x NA = 1.2 water objective) in a glass-bottomed 384-well plate that was pre-blocked with 1 mg/ml BSA. Saturation concentration was determined by quantifying the concentration in the dilute phase with a calibration curve acquired with concentration without condensation.

<u>Supported lipid bilayer (SLB) preparation</u>

*Small unilamellar vesicle (SUV) preparation*

Phospholipids including POPC (Avanti, Product No. 850457), certain amounts of DGS-NTA(Ni) (Avanti, Product No. 790404), 0.1 % PEG5000PE (Avanti, Product No. 880230) and 0.1 % Rhodamine-PE (Avanti, Product No. 810150) were mixed in chloroform in glass bottles, and dried under vacuum for 2 hrs. 500 uL to 1 mL buffer (20 mM Hepes, pH 7.4, 150 mM NaCl and 10 mM $MgCl_2$) was added to the dried lipid film to make a final lipid concentration as 1 mM and resuspended by shaking at a speed of 180 rpm, at 37 °C for 1 hr. The lipid mixture was then transferred to the Eppendorf tube and went through freeze-thaw for 15 runs until the lipid mixture became clear. The SUVs were further clarified by centrifugation at 17,000 g for 30 min and stored at 4 °C for use within one week.

*SLBs generation*

Glass-bottomed 96-well plates (Greiner, Product No. 655891) were pre-cleaned with 2 % Hellmanex II overnight and 6 M NaOH for 30 min at RT twice. Before adding SUVs, wells were equilibrated with buffers (20 mM Hepes, pH 7.4, 150 mM NaCl, and 10 mM $MgCl_2$) for 5 min, and left a 60 uL buffer in the well. 20 uL 1 mM SUVs were added and incubated for 20 min. 20 uL 5 M NaCl was then added for another 20 min for the SUVs to further collapse on the glass bottom. Excess SUVs were intensively washed away by pipetting in and out buffers (20 mM Hepes, pH 7.4, 150 mM NaCl, and 10 mM $MgCl_2$) 8 to 10 times. The quality of SLBs was checked by FRAP under a confocal microscope.

<u>ZO1 Surface phase separation assay on SLBs</u>

SLBs with a certain amount of DGS-NTA(Ni) lipids were blocked with 1 mg/ml BSA for 30 min. 500 nM monomeric, tetrameric, or 14-meric receptors were added and incubated for 30 min. Excess receptors were washed away by pipetting in and out buffer (20 mM Hepes, pH 7.4, 150 mM NaCl and 10 mM $MgCl_2$) 8 to 10 times. The images of receptors on SLBs before adding ZO1 were taken with a confocal microscope and converted to number density with respective calibration curves. The dynamics of receptors on SLBs before adding ZO1 were measured by FRAP. Certain amounts of ZO1-mGFP were added for 15 min to form ZO1 surface condensates. 3 images for each well were randomly taken under a confocal microscope. Time lapses were taken with a frame rate of 10 s for 15 min.

Client protein partition assay on SLBs

For the partition assay of cytosolic proteins including ZO2-mCherry, ZO3-mCherry, CGN-mCherry, AFDN-mCherry, PAR3-mCherry, and mCherry (used as a negative control), ZO1 surface condensates were formed by adding 100 nM ZO1 to membrane-bound 14-mer-DyLight 650 (680 molecules/$\mu m^2$) for 15 min. Images were taken after adding 100 nM client proteins to ZO1 surface condensates for 15 min. 3 images for each well were randomly taken under a confocal microscope.

For the partition assay of receptor proteins including CDH1-IC, JAM-A-IC, and CRB3-IC. 14-mer receptor-DyLight 650 (50 nM protein was used for the incubation) and client receptors (500 nM protein was used for the incubation) were anchored to SLBs containing 8 % DGS-NTA(Ni). Excess receptors were washed away by pipetting in and out buffer (20 mM Hepes, pH 7.4, 150 mM NaCl and 10 mM $MgCl_2$) 8 to 10 times. Receptors on SLBs before adding ZO1 were imaged with a confocal microscope and dynamically checked by FRAP. 100 nM ZO1 was added for 15 min to form ZO1 surface condensates.

Partition coefficients of client proteins into ZO1 surface condensates were quantified by measuring the fluorescence intensity of client proteins inside ZO1 surface condensates and normalized to the fluorescence intensity of client proteins outside ZO1 surface condensates.

Atomic force microscopy (AFM) imaging

For characterizing the height of ZO1 surface condensates, we performed AFM topography measurements at a JPK NanoWizard4 AFM (Bruker Nano GmbH, Germany). ZO1 surface condensates were prepared in 35 mm glass-bottomed dishes (MatTek Corporation, Product No. P35G-0.170-14-C). Images were acquired in JPK's "Q.I." mode (force mapping imaging) using "Biolever mini" cantilevers (BL-AC40TS-C2, Lot No. 840166, Olympus, Japan). Probes were calibrated before each measurement using the “contact-free” method implemented in the "NanoWizard Control Software" (version 6.1, JPK Instruments, Germany). The setpoints for the measurements were chosen between 0.1 and 0.2 nN, z-lengths between 100 and 250 nm, and pixel times between 6 and 10 ms.

Image processing was carried out using JPK's "Data Processing" software (version 6.1, JPK Instruments, Berlin, Germany). Maps of "height (measured)" were leveled line by line by fitting a first-order polynomial to the height values, only taking into account the substrate. Other than that, height maps were not subjected to further processing.

Muscle actin purification and labeling

Monomeric actin was purified from rabbit skeletal muscle acetone powder (Pel-freeze, USA, cat. no: 41995) following standard protocols (*48*) and kept at 4 °C throughout handling. A subset of depolymerized actin was labeled with either N-(1-Pyrenyl) iodoacetamide (Molecular Probes, USA) or Alexa647-maleimide (Thermo Fischer Scientific, cat. no: A20347). Therefore 2.5 mL of the dialyzed solution were buffer exchanged with a PD-10 column (Cytiva, USA) into DTT-free buffer (5 mM Tris-HCl pH 8, 0.2 mM ATP, 0.1 mM $CaCl_2$) and reacted at the equimolar ratio with Alexa-647 maleimide for 2 hrs in the dark. Aggregates have been removed by ultracentrifugation at 350 000 g for 20 min and actin in the supernatant was polymerized for 2 hrs by addition of 10x KMEI buffer (500 mM KCL, 10 mM $MgCl_2$, 10 mM EGTA, 100 mM Imidazole HCL, pH 7.0) and ATP to final concentrations 1x and 1 mM, respectively. Subsequently, actin filaments were pelleted again by ultracentrifugation at 100,000 g for 2 hrs, washed, and resuspended in 1.5 mL CaBuffer-G (2 mM Tris-HCl, pH 8.0, 0.2 mM ATP, 0.5 mM DTT, 1 mM NaN3, 0.1 mM $CaCl_2$) by homogenizing 10x with a douncer. The labeled actin solution was dialyzed against CaBuffer-G for 3 days to depolymerize actin. All actin solutions were filtered through 0.22 µm PTFE filters, ultracentrifuged for 2 hrs at 100,000 g, and gel-filtered on a superdex 200 column with CaBuffer-G. Only post-peak fractions were used for polymerization assays to assure a monomeric solution. Unlabeled and pyrene-labeled actin was stored at 4 °C for up to 4 weeks. Alexa647-labeled actin was flash-frozen with liquid nitrogen and stored at -80 °C. On the day of the experiment, Alexa647-labeled actin was rapidly thawed by hand and clarified by ultracentrifugation at 100,000 x g for 2 hrs at 4 °C using the top 75 % of the supernatant. Actin concentrations were determined in a nanophotometer (NP80, Implen, USA) at 290 nM (38.5 µM cm) and mixed to yield a 5 - 10 % labeled fraction.

Actin-pyrene polymerization assay in bulk

Actin assembly was monitored by pyrene fluorescence in 150 µL total reaction volume in 96 well formats with a Spark 20M (Tecan) plate reader (top reading, λex = 365 nm, λem = 407 nm, every 10 s, 90 min total time, T = 25 °C). Ca-ATP-actin (5 % pyrene-labeled) was converted to Mg-ATP-actin by incubation with 1/10$^{th}$ volume of 10x Mg exchange buffer (500 µM $MgCl_2$, 2 mM EGTA) for 2 min at RT. Mg-ATP actin was then mixed at a final concentration of 2 µM with the respective proteins (diluted to 1 µM in their buffer, final concentrations see Table 1 below) and transferred in up to 12 rows of a 96 well plate (half area, µClear, black, Greiner Bio-one, cat. no. 675090). Using a multichannel pipette Mg buffer-G (2 mM Tris-HCl, pH 8.0, 0.2 mM ATP, 0.5 mM DTT) and 1/10$^{th}$ volume of 10x KMEI buffer (500 mM KCl, 10 mM $MgCl_2$, 10 mM EGTA, 100 mM Imidazole-HCL, pH 7.0) were added to actin in all wells simultaneously and the recording was started within 10 s.

Table 1

| | C(start) in µM | C(final) in µM | V (uL) |
|---|---|---|---|
| Ca-ATP actin (5 % pyrene labeled) | 10 | 2 | 30.0 |
| 10x Mg exchange buffer | 10 | 1 | 3.0 |
| ZO1 | 2 | 0.2 | 15.0 |
| 10x KMEI buffer | 10 | 1 | 15.0 |
| Mg Buffer-G | | | 87 |
| V (total) | | | 150 |

Actin polymerization assay on SLBs

ZO1 surface condensates were prepared by adding 200 nM ZO1 to membranes functionalized with 14-mer-DyLight 550 or monomer-DyLight 550 receptors for 10 min in 50 µL total volume of actin polymerization buffer (2 mM Tris-HCl pH 8.0, 150 mM KCl, 10 mM $MgCl_2$, 0.2 mM ATP, 1 mM EGTA) supplemented with ATP regeneration and oxygen scavenger systems (*49*): phosphocreatine (10 mM), creatine phosphokinase (53 U/ml), pyranose oxidase (3.7 U/ml, Merck, cat. no. P4234), catalase (90 U/ml, Merck, cat. no. C40), 0.4 % glucose and 2mM 2-Mercaptoethanol.

Ca-ATP-actin (5 % AlexaFluor647-labeled) was converted to Mg-ATP-actin by incubation with 1/10th volume of 10x Mg exchange buffer (500 µM $MgCl_2$, 2 mM EGTA) for 2 min at RT. 15 ml of 13 µM Mg-ATP actin were then added to reach a final concentration of 3 µM and confocal time-lapse images were recorded with a frame rate of 30 s for 1 hr. Additionally, images at random spots were taken 1 h after the addition of actin.

For determining the dependence of actin polymerization on the receptor densities, membranes functionalized with different amounts of 14-mer receptors were used. Membranes without a receptor, without ZO1, or with ZO1-ΔABR were used as control. The length of ZO1 and actin networks were extracted with the plugin Skeleton in Fiji (https://fiji.sc/).

Production of F-actin visualization cell lines

CRISPR/Cas9 method was used to generate the N-terminal ZO1 NeonGreen (NG-ZO1) knock-in cell line in MDCK II cells as described before (*4*). On top of this, a mammalian expression plasmid with an N-terminal SNAP-tag and the first 261 amino acids of human utrophin, an F-actin binding protein, was transfected using Lipofectamine-2000. Stable transgenic cell lines were selected in the presence of geneticin (400 ug/mL). For visualizing the actin, SNAP-Utrophin stable expressing cell lines were labeled with 3 µM SNAP substrate (SiR-SNAP, Product No. S9102S) for 30 min before imaging. Free SNAP substrate was washed away before imaging.

Latrunculin-A (LatA) switch assay

ZO1-NG knock-in, SNAP-Utrophin stable expressing MDCK II cells were cultured in full medium (DMEM with 1 g/L glucose, supplemented with 5% FBS, 1% non-essential amino acids) at 37 ˚C with 5% $CO_2$ until they formed a confluent monolayer. For visualizing the actin, 3 µM SNAP substrates (SiR-SNAP, Product No. S9102S) were added for 30 min before imaging. Free SNAP substrate was washed away before imaging. 1 µM LatA was added for 30 min to disrupt the tight junction. LatA was then washed away. Time series images were taken under a Nikon spinning disk confocal every 80 s for 12 hrs to capture the elongation of tight junction belts after washing away the LatA.

Calcium switch assay

ZO1-NG knock-in, SNAP-Utrophin stable expressing MDCK II cells were cultured in full medium (DMEM with 1 g/L glucose, supplemented with 5% FBS, 1% non-essential amino acids) at 37 ˚C with 5% $CO_2$ until they formed a confluent monolayer. Confluent MDCK II monolayer was switched to a calcium depletion medium (Gibco, Product No. 21068) for 18 hrs until the tight junction was completely disrupted. For visualizing the actin, 3 µM SNAP substrates (SiR-SNAP, Product No. S9102S) were added for 30 min before imaging. Free SNAP substrate was washed away before imaging. The full medium with calcium was then switched back. Time series images were taken under a Nikon spinning disk confocal every 80 s for 12 hrs to capture the formation of tight junction belts.

**Supplement figures:**

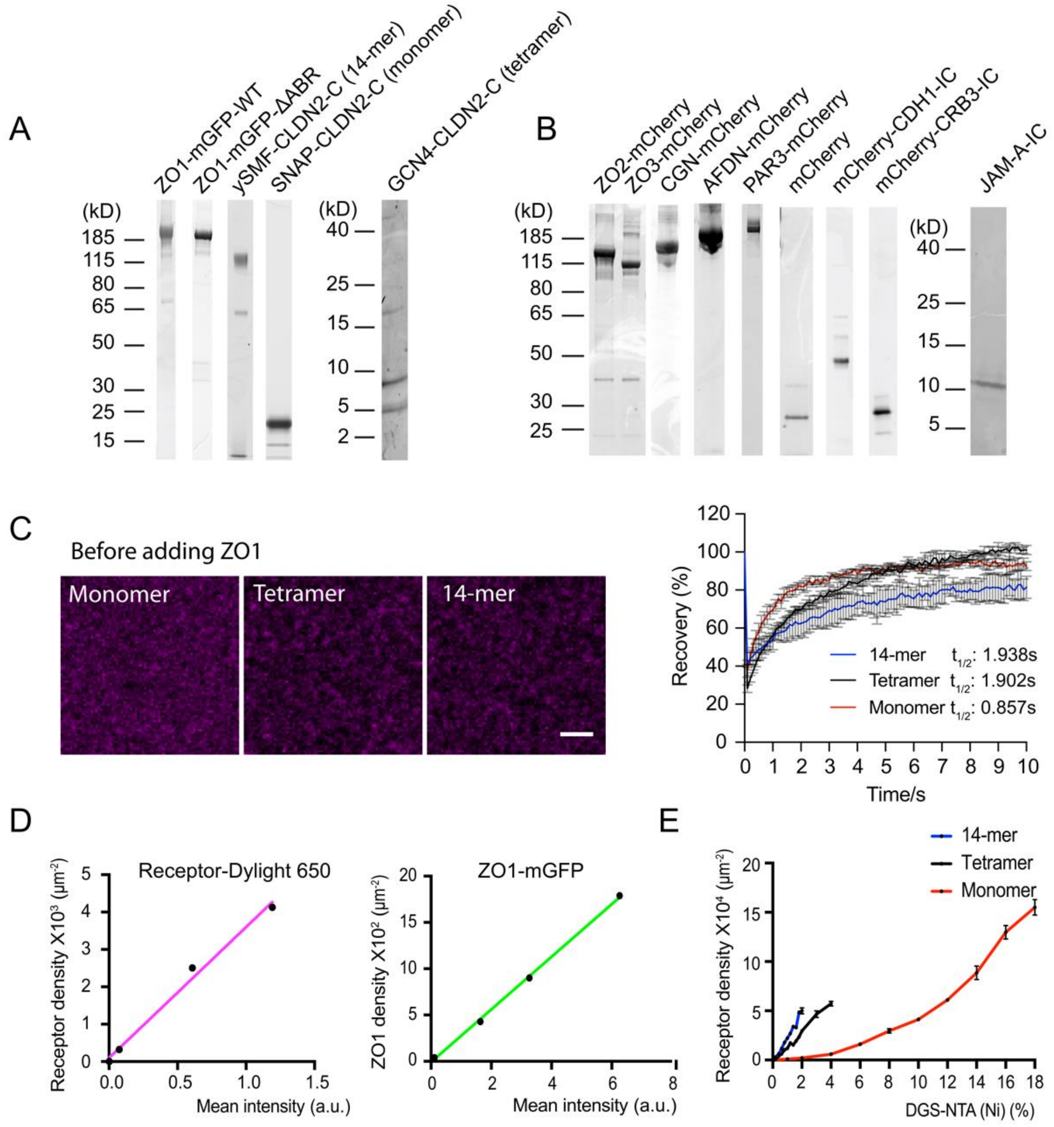

**Fig. S1. Receptors are dynamic before adding ZO1.** Related to Figure 1.

(A) Left: SDS-PAGE gel (stained with Coomassie blue) of ZO1-mGFP-WT, ZO1-mGFP-ΔABR, ySMF-CLDN2-C (14-meric receptor) and SNAP-CLDN2-C (monomeric receptor) proteins used in this study. Right: Tricine gel (stained with Coomassie blue) of GCN4-CLDN2-C (Tetrameric receptor) protein used in this study.

(B) Left: SDS-PAGE gel (stained with Coomassie blue) of mCherry tagged ZO2, ZO3, CGN, AFDN, PAR3, CDH1-IC, CRB3-IC and mCherry protein used in the partition assay. Right: Tricine gel (stained with Coomassie blue) of JAM-A-IC protein used in the partition assay.

(C) Left: Images of membranes functionalized with increasing amounts of monomeric, tetrameric, and 14-meric receptors. Receptors are all labeled with Dylight650. Right: Quantification of FRAP results of indicated receptors on membrane before adding ZO1. Values shown are the mean ± SD from 3 independent measurements. A one phase association fitting was performed to get the half-time of recovery. Half-times of recovery after photobleach are annotated.

(D) Left: Calibration curve of Dylight650 labeled receptors. The X-axis shows the mean fluorescence intensity acquired from confocal images; The Y-axis shows the molecules/$\mu m^2$ from FCS measurement. The calibration equation is Y = 3501*X + 97.29. Receptor numbers are in the unit of monomer. Right: Calibration curve of ZO1-mGFP. The X-axis shows the mean fluorescence intensity acquired from confocal images; The Y-axis shows the molecules/$\mu m^2$ from FCS measurement. The calibration equation is Y = 284.4*X – 8.990. ZO1-mGFP is in the unit of monomer.

(E) Quantification of indicated receptor densities with the increase of DGS-NTA(Ni) lipids on membranes. Receptor densities were converted to molecules/$\mu m^2$ from fluorescence intensity with the calibration curve in (D). Values shown are the mean ± SD from three different views.

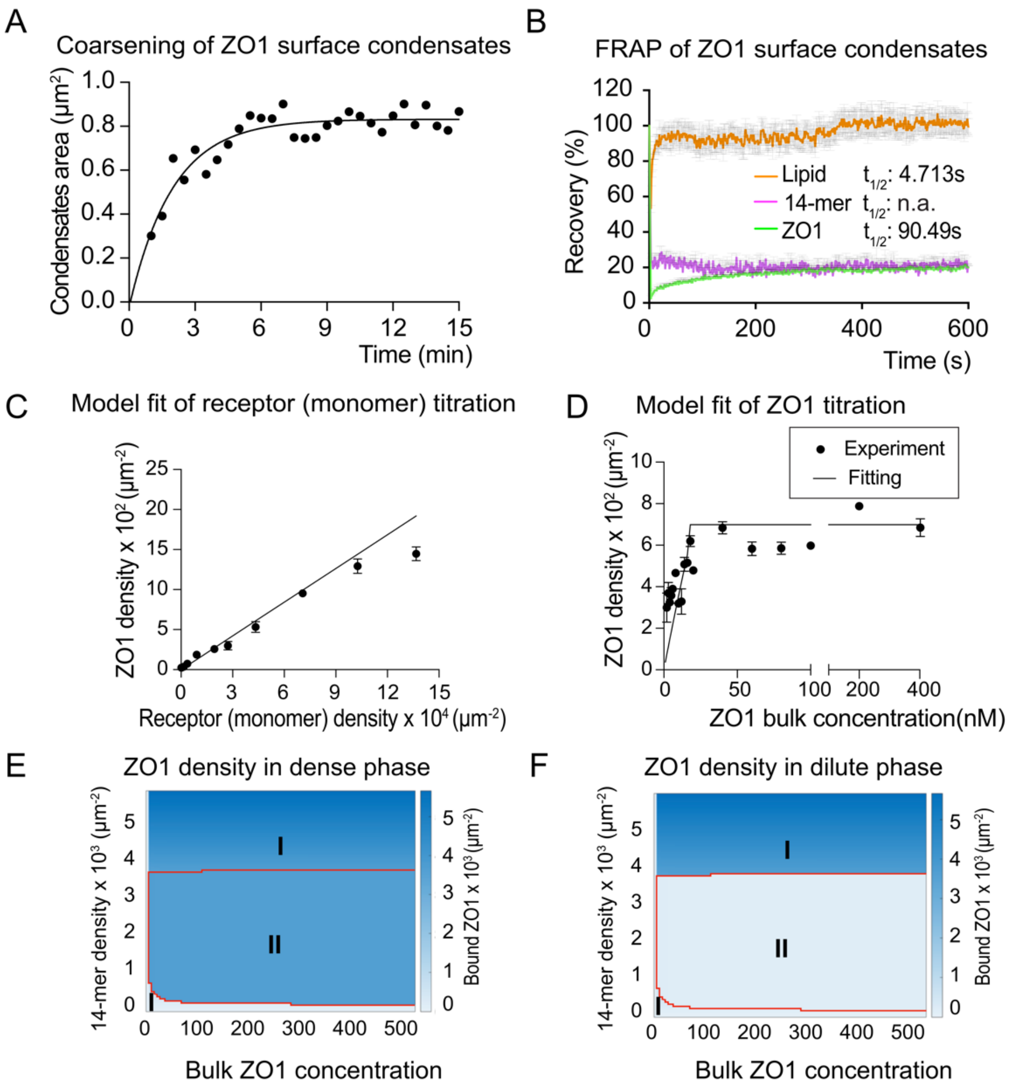

**Fig. S2. Fitting result of ZO1 membrane binding with monomer.** Related to Figure 2.

(A) Quantification of area change of ZO1 surface condensates after adding ZO1 from movie S1. A one phase association fitting was performed.

(B) Quantification of FRAP results of ZO1 surface condensates formed with 14-mer receptors. Values shown are the mean ± SD from 3 independent measurements. A one phase association fitting was performed to get the half-time of recovery. Half-times of recovery after photobleach are annotated.

(C) Quantification of ZO1 density on the membrane from simulation and experiment when titrating the monomer receptor densities. Values shown are the mean ± SD from three different views.

(D) Quantification of ZO1 density on the membrane from simulation and experiment when titrating the ZO1 bulk concentration with monomer receptors on the membrane. Values shown are the mean ± SD from three different views.

(E) Phase diagram of ZO1 density in dense phase with 14-mer receptors from simulation. Phase I shows the regime with mixed surface and mixed bulk. Phase II shows the regime with demixed surface and mixed bulk.

(F) Phase diagram of ZO1 density in dilute phase with 14-mer receptors from simulation. Phase I shows the regime with mixed surface and mixed bulk. Phase II shows the regime with demixed surface and mixed bulk.

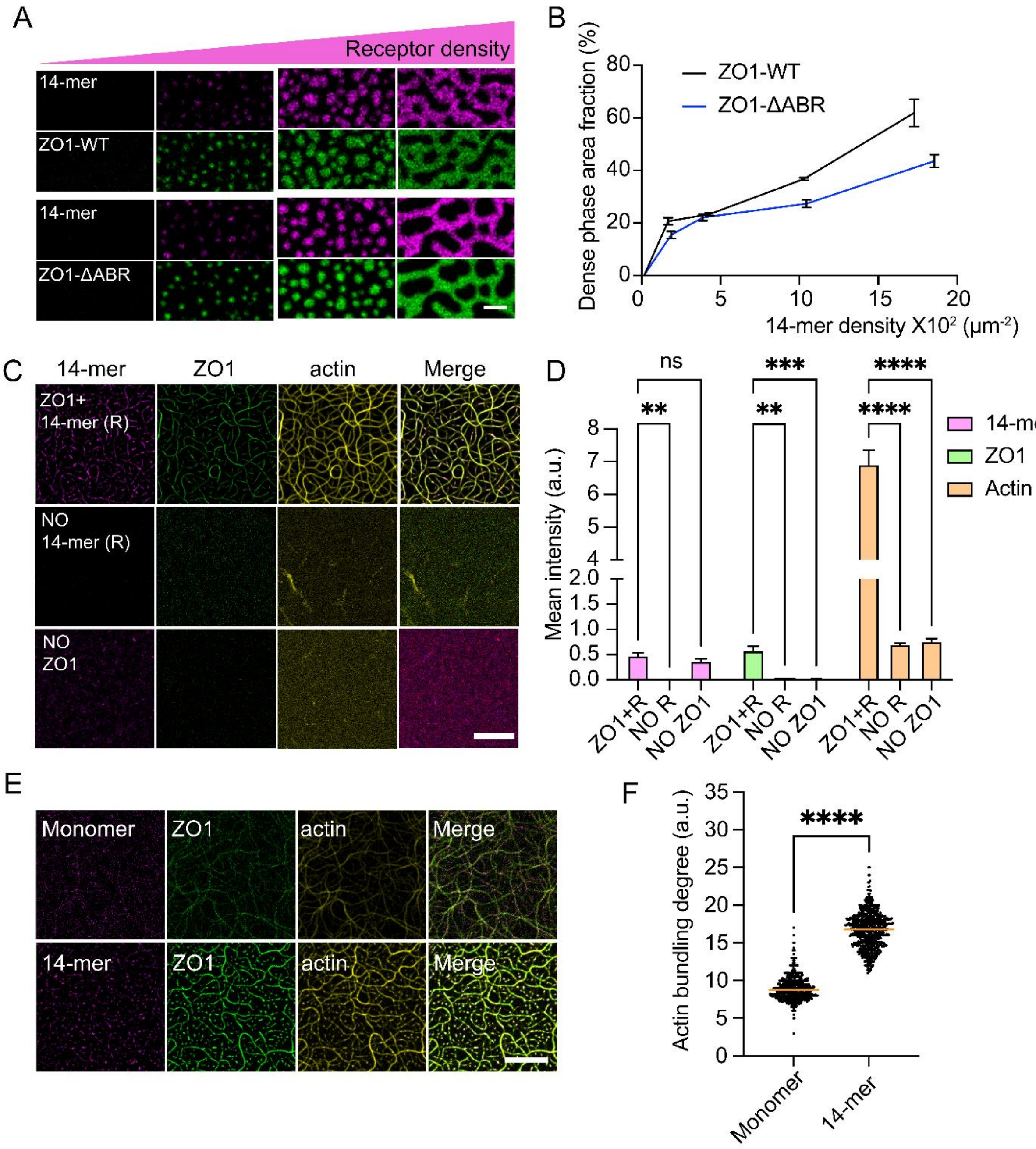

**Fig. S3. ZO1 surface condensates are vital for local actin polymerization.** Related to figure 4.

(A) Images of ZO1 surface condensates formed with ZO1-WT and ZO1-ΔABR. ZO1 surface condensates were formed by adding 200 nM ZO1-WT or ZO1-ΔABR to membranes functionalized with different amounts of 14-mer receptors. Scale bar, 2 μm.

(B) Dense phase area fraction quantification of ZO1 surface condensates from images in (A). Values shown are the mean ± SD from three different views.

(C) Actin local polymerization assay without ZO1 or 14-mer receptors. Scale bar, 10 μm.

(D) Quantification of ZO1, 14-mer receptor, and actin signal on the membrane from the images in (C). Values shown are the mean ± SD from 3 different views. A 2wayANOVA test was performed to determine the significance of the difference between different conditions.

(E) Actin polymerization assay with monomer or 14-mer on the membrane. 200 nM ZO1 protein was added to the membrane functionalized with_similar amounts of monomer (~3900 molecules/$\mu m^2$) or 14-mer (~260 molecules/$\mu m^2$) for 10 min. Images were taken at 60 min after adding 3 uM G-actin-Alexa Fluor 647. Scale bar, 10 µm.

(F) Actin bundling degree quantification from the images in (E). Values shown are the mean ± SD from three different views. An ordinary one-way ANOVA was performed to determine the significance of the difference between monomer and 14-mer.

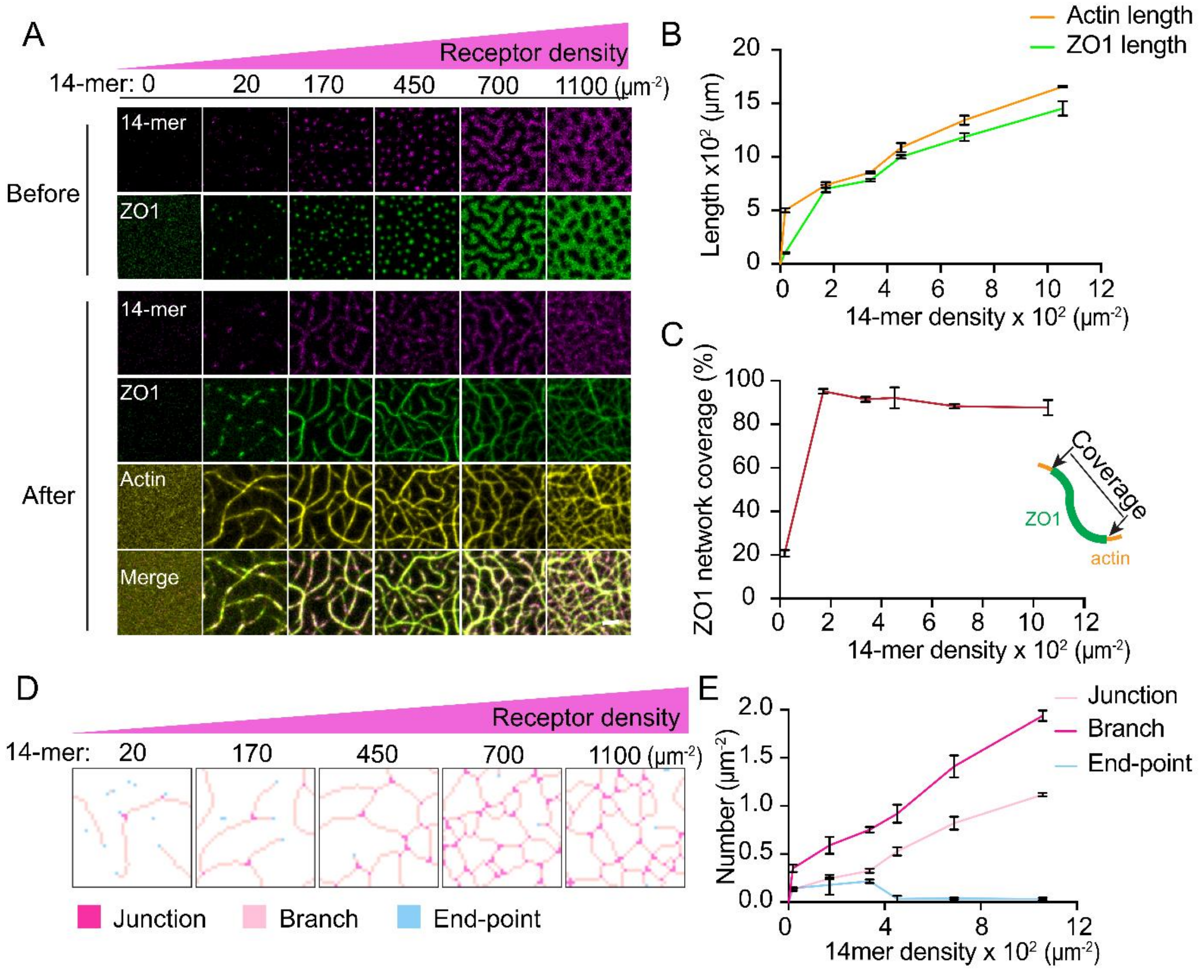

**Fig. S4. Tight junction continuity depends on receptor density.** Related to Figure 4.

(A) Before: Images of ZO1 surface condensates formed with 14-mer receptors. ZO1 membrane condensates were formed by adding 200 nM ZO1 protein to membranes functionalized with different amounts of 14-mer-Dylight 550 receptors. 14-mer receptor densities were annotated on the top. After: Images of tight junction like networks after adding 3 uM G-actin-Alexa Fluor 647 to ZO1 membrane condensates for 3 hrs. Scale bar, 2 μm.

(B) Network length quantification from ZO1 and actin signal after adding 3 uM G-actin-Alexa Fluor 647 to ZO1 surface condensates for 3 hrs from images in (A). Values shown are the mean ± SD from three different views.

(C) Quantification of coverage degree of ZO1 network on actin networks from (B). Values shown are the mean ± SD from three different views.

(D) Network complexity analysis by skeletonizing actin networks from the images in (A) with Skeletonize Plugin in FIJI (https://imagej.net/plugins/skeletonize3d). The skeleton is classified into end-point (less than 2 neighbors), branch (exactly 2 neighbors) and junction (more than 2 neighbors) by the number of neighbors.

(E) Quantification of network complexity from the skeletonized actin network in (D). Values shown are the mean ± SD from three different views.

**Table S1. Sequences of constructs used in this study.**

| Construct | Sequence | Notes |
|---|---|---|
| ZO1-mGFP-WT | MGHHHHHHSSGRMKIEEGKLVIWINGDKGYNGLAEVG<br>KKFEKDTGIKVTVEHPDKLEEKFPQVAATGDGPDIIFWA<br>HDRFGGYAQSGLLAEITPDKAFQDKLYPFTWDAVRYN<br>GKLIAYPIAVEALSLIYNKDLLPNPPKTWEEIPALDKELK<br>AKGKSALMFNLQEPYFTWPLIAADGGYAFKYENGKYDI<br>KDVGVDNAGAKAGLTFLVDLIKNKHMNADTDYSIAEA<br>AFNKGETAMTINGPWAWSNIDTSKVNYGVTVLPTFKG<br>QPSKPFVGVLSAGINAASPNKELAKEFLENYLLTDEGLE<br>AVNKDKPLGAVALKSYEEELVKDPRIAATMENAQKGEI<br>MPNIPQMSAFWYAVRTAVINAASGRQTVDEALKDAQT<br>NSSSNNNNNNNNNNSSGRLEVLFQGPAAAMSARAAAA<br>KSTAMEETAIWEQHTVTLHRAPGFGFGIAISGGRDNPHF<br>QSGETSIVISDVLKGGPAEGQLQENDRVAMVNGVSMD<br>NVEHAFAVQQLRKSGKNAKITIRRKKKVQIPVSRPDPEP<br>VSDNEEDSYDEEIHDPRSGRSGVVNRRSEKIWPRDRSAS<br>RERSLSPRSDRRSVASSQPAKPTKVTLVKSRKNEEYGLR<br>LASHIFVKEISQDSLAARDGNIQEGDVVLKINGTVTENM<br>SLTDAKTLIERSKGKLKMVVQRDERATLLNVPDLSDSIH<br>SANASERDDISEIQSLASDHSGRSHDRPPRRSRSRSPDQR<br>SEPSDHSRHSPQQPSNGSLRSRDEERISKPGAVSTPVKHA<br>DDHTPKTVEEVTVERNEKQTPSLPEPKPVYAQVGQPDV<br>DLPVSPSDGVLPNSTHEDGILRPSMKLVKFRKGDSVGLR<br>LAGGNDVGIFVAGVLEDSPAAKEGLEEGDQILRVNNVD<br>FTNIIREEAVLFLLDLPKGEEVTILAQKKKDVYRRIVESD<br>VGDSFYIRTHFEYEKESPYGLSFNKGEVFRVVDTLYNG<br>KLGSWLAIRIGKNHKEVERGIIPNKNRAEQLASVQYTLP<br>KTAGGDRADFWRFRGLRSSKRNLRKSREDLSAQPVQT<br>KFPAYERVVLREAGFLRPVTIFGPIADVAREKLAREEPDI<br>YQIAKSEPRDAGTDQRSSGIIRLHTIKQIIDQDKHALLDV<br>TPNAVDRLNYAQWYPIVVFLNPDSKQGVKTMRMRLCP<br>ESRKSARKLYERSHKLRKNNHHLFTTTINLNSMNDGWY<br>GALKEAIQQQQNQLVWVSEGKADGATSDDLDLHDDRL<br>SYLSAPGSEYSMYSTDSRHTSDYEDTDTEGGAYTDQEL<br>DETLNDEVGTPPESAITRSSEPVREDSSGMHHENQTYPP<br>YSPQAQPQPIHRIDSPGFKPASQQKAEASSPVPYLSPETN<br>PASSTSAVNHNVNLTNVRLEEPTPAPSTSYSPQADSLRT<br>PSTEAAHIMLRDQEPSLSSHVDPTKVYRKDPYPEEMMR | Full length human ZO1, fused with an N-terminal His6-MBP tag and C-terminal mGFP tag |

| | | |
|---|---|---|
| | QNHVLKQPAVSHPGHRPDKEPNLTYEPQLPYVEKQASR<br>DLEQPTYRYESSSYTDQFSRNYEHRLRYEDRVPMYEEQ<br>WSYYDDKQPYPSRPPFDNQHSQDLDSRQHPEESSERGY<br>FPRFEEPAPLSYDSRPRYEQAPRASALRHEEQPAPGYDT<br>HGRLRPEAQPHPSAGPKPAESKQYFEQYSRSYEQVPPQ<br>GFTSRAGHFEPLHGAAAVPPLIPSSQHKPEALPSNTKPLP<br>PPPTQTEEEEDPAMKPQSVLTRVKMFENKRSASLETKK<br>DVNDTGSFKPPEVASKPSGAPIIGPKPTSQNQFSEHDKTL<br>YRIPEPQKPQLKPPEDIVRSNHYDPEEDEEYYRKQLSYF<br>DRRSFENKPPAHIAASHLSEPAKPAHSQNQSNFSSYSSK<br>GKPPEADGVDRSFGEKRYEPIQATPPPPPLPSQYAQPSQP<br>VTSASLHIHSKGAHGEGNSVSLDFQNSLVSKPDPPPSQN<br>KPATFRPPNREDTAQAAFYPQKSFPDKAPVNGTEQTQK<br>TVTPAYNRFTPKPYTSSARPFERKFESPKFNHNLLPSETA<br>HKPDLSSKTPTSPKTLVKSHSLAQPPEFDSGVETFSIHAE<br>KPKYQINNISTVPKAIPVSPSAVEEDEDEDGHTVVATAR<br>GIFNSNGGVLSSIETGVSIIIPQGAIPEGVEQEIYFKVCRD<br>NSILPPLDKEKGETLLSPLVMCGPHGLKFLKPVELRLPH<br>CDPKTWQNKCLPGDPNYLVGANCVSVLIDHFGAPGSSS<br>GRENLYFQGMVSKGEELFTGVVPILVELDGDVNGHKFS<br>VSGEGEGDATYGKLTLKFICTTGKLPVPWPTLVTTLTY<br>GVQCFSRYPDHMKQHDFFKSAMPEGYVQERTIFFKDDG<br>NYKTRAEVKFEGDTLVNRIELKGIDFKEDGNILGHKLEY<br>NYNSHNVYIMADKQKNGIKVNFKIRHNIEDGSVQLADH<br>YQQNTPIGDGPVLLPDNHYLSTQSKLSKDPNEKRDHMV<br>LLEFVTAAGITLGMDELYK | |
| ZO1-<br>mCherry | MGMKIEEGKLVIWINGDKGYNGLAEVGKKFEKDTGIK<br>VTVEHPDKLEEKFPQVAATGDGPDIIFWAHDRFGGYAQ<br>SGLLAEITPDKAFQDKLYPFTWDAVRYNGKLIAYPIAVE<br>ALSLIYNKDLLPNPPKTWEEIPALDKELKAKGKSALMFN<br>LQEPYFTWPLIAADGGYAFKYENGKYDIKDVGVDNAG<br>AKAGLTFLVDLIKNKHMNADTDYSIAEAAFNKGETAM<br>TINGPWAWSNIDTSKVNYGVTVLPTFKGQPSKPFVGVL<br>SAGINAASPNKELAKEFLENYLLTDEGLEAVNKDKPLG<br>AVALKSYEEELVKDPRIAATMENAQKGEIMPNIPQMSA<br>FWYAVRTAVINAASGRQTVDEALKDAQTNSSSNNNNN<br>NNNNNSSGRLEVLFQGPAAAMSARAAAAKSTAMEETA<br>IWEQHTVTLHRAPGFGFGIAISGGRDNPHFQSGETSIVIS<br>DVLKGGPAEGQLQENDRVAMVNGVSMDNVEHAFAVQ | Full length human ZO1, fused with an N-terminal MBP tag and C-terminal mCherry-His6 tag |

| | |
|---|---|
| | QLRKSGKNAKITIRRKKKVQIPVSRPDPEPVSDNEEDSY<br>DEEIHDPRSGRSGVVNRRSEKIWPRDRSASRERSLSPRS<br>DRRSVASSQPAKPTKVTLVKSRKNEEYGLRLASHIFVKE<br>ISQDSLAARDGNIQEGDVVLKINGTVTENMSLTDAKTLI<br>ERSKGKLKMVVQRDERATLLNVPDLSDSIHSANASERD<br>DISEIQSLASDHSGRSHDRPPRRSRSRSPDQRSEPSDHSR<br>HSPQQPSNGSLRSRDEERISKPGAVSTPVKHADDHTPKT<br>VEEVTVERNEKQTPSLPEPKPVYAQVGQPDVDLPVSPS<br>DGVLPNSTHEDGILRPSMKLVKFRKGDSVGLRLAGGND<br>VGIFVAGVLEDSPAAKEGLEEGDQILRVNNVDFTNIIRE<br>EAVLFLLDLPKGEEVTILAQKKKDVYRRIVESDVGDSFY<br>IRTHFEYEKESPYGLSFNKGEVFRVVDTLYNGKLGSWL<br>AIRIGKNHKEVERGIIPNKNRAEQLASVQYTLPKTAGGD<br>RADFWRFRGLRSSKRNLRKSREDLSAQPVQTKFPAYER<br>VVLREAGFLRPVTIFGPIADVAREKLAREEPDIYQIAKSE<br>PRDAGTDQRSSGIIRLHTIKQIIDQDKHALLDVTPNAVDR<br>LNYAQWYPIVVFLNPDSKQGVKTMRMRLCPESRKSAR<br>KLYERSHKLRKNNHHLFTTTINLNSMNDGWYGALKEAI<br>QQQQNQLVWVSEGKADGATSDDLDLHDDRLSYLSAPG<br>SEYSMYSTDSRHTSDYEDTDTEGGAYTDQELDETLNDE<br>VGTPPESAITRSSEPVREDSSGMHHENQTYPPYSPQAQP<br>QPIHRIDSPGFKPASQQKAEASSPVPYLSPETNPASSTSA<br>VNHNVNLTNVRLEEPTPAPSTSYSPQADSLRTPSTEAAH<br>IMLRDQEPSLSSHVDPTKVYRKDPYPEEMMRQNHVLK<br>QPAVSHPGHRPDKEPNLTYEPQLPYVEKQASRDLEQPT<br>YRYESSSYTDQFSRNYEHRLRYEDRVPMYEEQWSYYD<br>DKQPYPSRPPFDNQHSQDLDSRQHPEESSERGYFPRFEE<br>PAPLSYDSRPRYEQAPRASALRHEEQPAPGYDTHGRLRP<br>EAQPHPSAGPKPAESKQYFEQYSRSYEQVPPQGFTSRAG<br>HFEPLHGAAAVPPLIPSSQHKPEALPSNTKPLPPPPTQTE<br>EEEDPAMKPQSVLTRVKMFENKRSASLETKKDVNDTGS<br>FKPPEVASKPSGAPIIGPKPTSQNQFSEHDKTLYRIPEPQK<br>PQLKPPEDIVRSNHYDPEEDEEYYRKQLSYFDRRSFENK<br>PPAHIAASHLSEPAKPAHSQNQSNFSSYSSKGKPPEADG<br>VDRSFGEKRYEPIQATPPPPPLPSQYAQPSQPVTSASLHI<br>HSKGAHGEGNSVSLDFQNSLVSKPDPPPSQNKPATFRPP<br>NREDTAQAAFYPQKSFPDKAPVNGTEQTQKTVTPAYNR<br>FTPKPYTSSARPFERKFESPKFNHNLLPSETAHKPDLSSK<br>TPTSPKTLVKSHSLAQPPEFDSGVETFSIHAEKPKYQINN<br>ISTVPKAIPVSPSAVEEDEDEDGHTVVATARGIFNSNGG |

| | | |
|---|---|---|
| | VLSSIETGVSIIIPQGAIPEGVEQEIYFKVCRDNSILPPLDK<br>EKGETLLSPLVMCGPHGLKFLKPVELRLPHCDPKTWQN<br>KCLPGDPNYLVGANCVSVLIDHFGAPGSAGSAAGSGM<br>VSKGEEDNMAIIKEFMRFKVHMEGSVNGHEFEIEGEGE<br>GRPYEGTQTAKLKVTKGGPLPFAWDILSPQFMYGSKAY<br>VKHPADIPDYLKLSFPEGFKWERVMNFEDGGVVTVTQ<br>DSSLQDGEFIYKVKLRGTNFPSDGPVMQKKTMGWEASS<br>ERMYPEDGALKGEIKQRLKLKDGGHYDAEVKTTYKAK<br>KPVQLPGAYNVNIKLDITSHNEDYTIVEQYERAEGRHST<br>GGMDELYKLEVLFQGPGSSHHHHHHSG | |
| ZO1-mGFP-<br>ΔABR | MGMKIEEGKLVIWINGDKGYNGLAEVGKKFEKDTGIK<br>VTVEHPDKLEEKFPQVAATGDGPDIIFWAHDRFGGYAQ<br>SGLLAEITPDKAFQDKLYPFTWDAVRYNGKLIAYPIAVE<br>ALSLIYNKDLLPNPPKTWEEIPALDKELKAKGKSALMFN<br>LQEPYFTWPLIAADGGYAFKYENGKYDIKDVGVDNAG<br>AKAGLTFLVDLIKNKHMNADTDYSIAEAAFNKGETAM<br>TINGPWAWSNIDTSKVNYGVTVLPTFKGQPSKPFVGVL<br>SAGINAASPNKELAKEFLENYLLTDEGLEAVNKDKPLG<br>AVALKSYEEELVKDPRIAATMENAQKGEIMPNIPQMSA<br>FWYAVRTAVINAASGRQTVDEALKDAQTNSSSNNNNN<br>NNNNNSSGRLEVLFQGPAAAMSARAAAAKSTAMEETA<br>IWEQHTVTLHRAPGFGFGIAISGGRDNPHFQSGETSIVIS<br>DVLKGGPAEGQLQENDRVAMVNGVSMDNVEHAFAVQ<br>QLRKSGKNAKITIRRKKKVQIPVSRPDPEPVSDNEEDSY<br>DEEIHDPRSGRSGVVNRRSEKIWPRDRSASRERSLSPRS<br>DRRSVASSQPAKPTKVTLVKSRKNEEYGLRLASHIFVKE<br>ISQDSLAARDGNIQEGDVVLKINGTVTENMSLTDAKTLI<br>ERSKGKLKMVVQRDERATLLNVPDLSDSIHSANASERD<br>DISEIQSLASDHSGRSHDRPPRRSRSRSPDQRSEPSDHSR<br>HSPQQPSNGSLRSRDEERISKPGAVSTPVKHADDHTPKT<br>VEEVTVERNEKQTPSLPEPKPVYAQVGQPDVDLPVSPS<br>DGVLPNSTHEDGILRPSMKLVKFRKGDSVGLRLAGGND<br>VGIFVAGVLEDSPAAKEGLEEGDQILRVNNVDFTNIIRE<br>EAVLFLLDLPKGEEVTILAQKKKDVYRRIVESDVGDSFY<br>IRTHFEYEKESPYGLSFNKGEVFRVVDTLYNGKLGSWL<br>AIRIGKNHKEVERGIIPNKNRAEQLASVQYTLPKTAGGD<br>RADFWRFRGLRSSKRNLRKSREDLSAQPVQTKFPAYER<br>VVLREAGFLRPVTIFGPIADVAREKLAREEPDIYQIAKSE<br>PRDAGTDQRSSGIIRLHTIKQIIDQDKHALLDVTPNAVDR | Human ZO1 without actin binding region (Δ1152-1375), fused with a N-terminal MBP tag and C-terminal mGFP-His6 tag |

| | | |
|---|---|---|
| | LNYAQWYPIVVFLNPDSKQGVKTMRMRLCPESRKSAR KLYERSHKLRKNNHHLFTTTINLNSMNDGWYGALKEAI QQQQNQLVWVSEGKADGATSDDLDLHDDRLSYLSAPG SEYSMYSTDSRHTSDYEDTDTEGGAYTDQELDETLNDE VGTPPESAITRSSEPVREDSSGMHHENQTYPPYSPQAQP QPIHRIDSPGFKPASQQKAEASSPVPYLSPETNPASSTSA VNHNVNLTNVRLEEPTPAPSTSYSPQADSLRTPSTEAAH IMLRDQEPSLSSHVDPTKVYRKDPYPEEMMRQNHVLK QPAVSHPGHRPDKEPNLTYEPQLPYVEKQASRDLEQPT YRYESSSYTDQFSRNYEHRLRYEDRVPMYEEQWSYYD DKQPYPSRPPFDNQHSQDLDSRQHPEESSERGYFPRFEE PAPLSYDSRPRYEQAPRAAASHLSEPAKPAHSQNQSNFS SYSSKGKPPEADGVDRSFGEKRYEPIQATPPPPPLPSQYA QPSQPVTSASLHIHSKGAHGEGNSVSLDFQNSLVSKPDP PPSQNKPATFRPPNREDTAQAAFYPQKSFPDKAPVNGTE QTQKTVTPAYNRFTPKPYTSSARPFERKFESPKFNHNLL PSETAHKPDLSSKTPTSPKTLVKSHSLAQPPEFDSGVETF SIHAEKPKYQINNISTVPKAIPVSPSAVEEDEDEDGHTVV ATARGIFNSNGGVLSSIETGVSIIIPQGAIPEGVEQEIYFK VCRDNSILPPLDKEKGETLLSPLVMCGPHGLKFLKPVEL RLPHCDPKTWQNKCLPGDPNYLVGANCVSVLIDHFGAP GSSSGRENLYFQGMVSKGEELFTGVVPILVELDGDVNG HKFSVSGEGEGDATYGKLTLKFICTTGKLPVPWPTLVTT LTYGVQCFSRYPDHMKQHDFFKSAMPEGYVQERTIFFK DDGNYKTRAEVKFEGDTLVNRIELKGIDFKEDGNILGH KLEYNYNSHNVYIMADKQKNGIKVNFKIRHNIEDGSVQ LADHYQQNTPIGDGPVLLPDNHYLSTQSKLSKDPNEKR DHMVLLEFVTAAGITLGMDELYKLEVLFQGPGSSHHHH HHSG | |
| SNAP-CLDN2-C7 (monomer) | MGSSHHHHHHHHHHSSGRLEVLFQGPMDKDCEMKRTT LDSPLGKLELSGCEQGLHRIIFLGKGTSAADAVEVPAPA AVLGGPEPLMQATAWLNAYFHQPEAIEEFPVPALHHPV FQQESFTRQVLWKLLKVVKFGEVISYSHLAALAGNPAA TAAVKTALSGNPVPILIPCHRVVQGDLDVGGYEGGLAV KEWLLAHEGHRLGKPGLGGSAGSAAGSGAAAYSLTGY V | Last 7 amino acids from human CLDN2, fused with an N-terminal His10-SNAP tag, referred as monomer in this study |

| | | |
|---|---|---|
| GCN4-CLDN2-C7 (tetramer) | MGSSHHHHHHSSGRENLYFQGGGDSLEFIASKLAGSGSGSAAAMKQIEDKLEEILSKLYHIENELARIKKLLGERYSLTGYV | Last 7 amino acids from human CLDN2, fused with an His6 tag and a GCN4 domain on the N-terminus, referred as tetramer in this study |
| ySMF-CLDN2-C7 (14-mer) | MGSSHHHHHHSSGRENLYFQGGGDSLEFIASKLAGSGSGSAAAMSESSDISAMQPVNPKPFLKGLVNHRVGVKLKFNSTEYRGTLVSTDNYFNLQLNEAEEFVAGVSHGTLGEIFIRSNNVLYIRELPNYSLTGYV | Last 7 amino acids from human CLDN2, fused with an His6 tag and a ySMF domain on the N-terminus, referred as 14-mer in this study |
| mCherry-ZO2 | MGMKIEEGKLVIWINGDKGYNGLAEVGKKFEKDTGIKVTVEHPDKLEEKFPQVAATGDGPDIIFWAHDRFGGYAQSGLLAEITPDKAFQDKLYPFTWDAVRYNGKLIAYPIAVEALSLIYNKDLLPNPPKTWEEIPALDKELKAKGKSALMFNLQEPYFTWPLIAADGGYAFKYENGKYDIKDVGVDNAGAKAGLTFLVDLIKNKHMNADTDYSIAEAAFNKGETAMTINGPWAWSNIDTSKVNYGVTVLPTFKGQPSKPFVGVLSAGINAASPNKELAKEFLENYLLTDEGLEAVNKDKPLGAVALKSYEEELVKDPRIAATMENAQKGEIMPNIPQMSAFWYAVRTAVINAASGRQTVDEALKDAQTNSSSNNNNNNNNNNSSGRLEVLFQGPAAAMPVRGDRGFPPRRELSGWLRAPGMEELIWEQYTVTLQKDSKRGFGIAVSGGRDNPHFENGETSIVISDVLPGGPADGLLQENDRVVMVNGTPMEDVLHSFAVQQLRKSGKVAAIVVKRPRKVQVAALQAS | Full length human ZO2, fused with an N-terminal MBP tag and C-terminal mCherry-His6 tag |

| | |
|---|---|
| | PPLDQDDRAFEVMDEFDGRSFRSGYSERSRLNSHGGRS<br>RSWEDSPERGRPHERARSRERDLSRDRSRGRSLERGLD<br>QDHARTRDRSRGRSLERGLDHDFGPSRDRDRDRSRGRS<br>IDQDYERAYHRAYDPDYERAYSPEYRRGARHDARSRG<br>PRSRSREHPHSRSPSPEPRGRPGPIGVLLMKSRANEEYGL<br>RLGSQIFVKEMTRTGLATKDGNLHEGDIILKINGTVTEN<br>MSLTDARKLIEKSRGKLQLVVLRDSQQTLINIPSLNDSD<br>SEIEDISEIESNRSFSPEERRHQYSDYDYHSSSEKLKERPS<br>SREDTPSRLSRMGATPTPFKSTGDIAGTVVPETNKEPRY<br>QEDPPAPQPKAAPRTFLRPSPEDEAIYGPNTKMVRFKKG<br>DSVGLRLAGGNDVGIFVAGIQEGTSAEQEGLQEGDQIL<br>KVNTQDFRGLVREDAVLYLLEIPKGEMVTILAQSRADV<br>YRDILACGRGDSFFIRSHFECEKETPQSLAFTRGEVFRVV<br>DTLYDGKLGNWLAVRIGNELEKGLIPNKSRAEQMASV<br>QNAQRDNAGDRADFWRMRGQRSGVKKNLRKSREDLT<br>AVVSVSTKFPAYERVLLREAGFKRPVVLFGPIADIAMEK<br>LANELPDWFQTAKTEPKDAGSEKSTGVVRLNTVRQIIE<br>QDKHALLDVTPKAVDLLNYTQWFPIVIFFNPDSRQGVK<br>TMRQRLNPTSNKSSRKLFDQANKLKKTCAHLFTATINL<br>NSANDSWFGSLKDTIQHQQGEAVWVSEGKMEGMDDD<br>PEDRMSYLTAMGADYLSCDSRLISDFEDTDGEGGAYTD<br>NELDEPAEEPLVSSITRSSEPVQHEESIRKPSPEPRAQMR<br>RAASSDQLRDNSPPPAFKPEPPKAKTQNKEESYDFSKSY<br>EYKSNPSAVAGNETPGASTKGYPPPVAAKPTFGRSILKP<br>STPIPPQEGEEVGESSEEQDNAPKSVLGKVKIFEKMDHK<br>ARLQRMQELQEAQNARIEIAQKHPDIYAVPIKTHKPDPG<br>TPQHTSSRPPEPQKAPSRPYQDTRGSYGSDAEEEEYRQQ<br>LSEHSKRGYYGQSARYRDTELGAPGSAGSAAGSGMVS<br>KGEEDNMAIIKEFMRFKVHMEGSVNGHEFEIEGEGEGR<br>PYEGTQTAKLKVTKGGPLPFAWDILSPQFMYGSKAYVK<br>HPADIPDYLKLSFPEGFKWERVMNFEDGGVVTVTQDSS<br>LQDGEFIYKVKLRGTNFPSDGPVMQKKTMGWEASSER<br>MYPEDGALKGEIKQRLKLKDGGHYDAEVKTTYKAKKP<br>VQLPGAYNVNIKLDITSHNEDYTIVEQYERAEGRHSTGG<br>MDELYKLEVLFQGPGSSHHHHHHSG |

| | | |
|---|---|---|
| mCherry-<br>ZO3 | MGMKIEEGKLVIWINGDKGYNGLAEVGKKFEKDTGIK<br>VTVEHPDKLEEKFPQVAATGDGPDIIFWAHDRFGGYAQ<br>SGLLAEITPDKAFQDKLYPFTWDAVRYNGKLIAYPIAVE<br>ALSLIYNKDLLPNPPKTWEEIPALDKELKAKGKSALMFN<br>LQEPYFTWPLIAADGGYAFKYENGKYDIKDVGVDNAG<br>AKAGLTFLVDLIKNKHMNADTDYSIAEAAFNKGETAM<br>TINGPWAWSNIDTSKVNYGVTVLPTFKGQPSKPFVGVL<br>SAGINAASPNKELAKEFLENYLLTDEGLEAVNKDKPLG<br>AVALKSYEEELVKDPRIAATMENAQKGEIMPNIPQMSA<br>FWYAVRTAVINAASGRQTVDEALKDAQTNSSSNNNNN<br>NNNNNSSGRLEVLFQGPAAAMEELTIWEQHTATLSKDP<br>RRGFGIAISGGRDRPGGSMVVSDVVPGGPAEGRLQTGD<br>HIVMVNGVSMENATSAFAIQILKTCTKMANITVKRPRRI<br>HLPATKASPSSPGRQDSDEDDGPQRVEEVDQGRGYDGD<br>SSSGSGRSWDERSRRPRPGRRGRAGSHGRRSPGGGSEA<br>NGLALVSGFKRLPRQDVQMKPVKSVLVKRRDSEEFGV<br>KLGSQIFIKHITDSGLAARHRGLQEGDLILQINGVSSQNL<br>SLNDTRRLIEKSEGKLSLLVLRDRGQFLVNIPPAVSDSDS<br>SPLEDISDLASELSQAPPSHIPPPPRHAQRSPEASQTDSPV<br>ESPRLRRESSVDSRTISEPDEQRSELPRESSYDIYRVPSSQ<br>SMEDRGYSPDTRVVRFLKGKSIGLRLAGGNDVGIFVSG<br>VQAGSPADGQGIQEGDQILQVNDVPFQNLTREEAVQFL<br>LGLPPGEEMELVTQRKQDIFWKMVQSRVGDSFYIRTHF<br>ELEPSPPSGLGFTRGDVFHVLDTLHPGPGQSHARGGHW<br>LAVRMGRDLREQERGIIPNQSRAEQLASLEAAQRAVGV<br>GPGSSAGSNARAEFWRLRGLRRGAKKTTQRSREDLSAL<br>TRQGRYPPYERVVLREASFKRPVVILGPVADIAMQKLT<br>AEMPDQFEIAETVSRTDSPSKIIKLDTVRVIAEKDKHALL<br>DVTPSAIERLNYVQYYPIVVFFIPESRPALKALRQWLAP<br>ASRRSTRRLYAQAQKLRKHSSHLFTATIPLNGTSDTWY<br>QELKAIIREQQTRPIWTAEDQLDGSLEDNLDLPHHGLAD<br>SSADLSCDSRVNSDYETDGEGGAYTDGEGYTDGEGGP<br>YTDVDDEPPAPALARSSEPVQADESQSPRDRGRISAHQG<br>AQVDSRHPQGQWRQDSMRTYEREALKKKFMRVHDAE<br>SSDEDGYDWGPATDLGAPGSAGSAAGSGMVSKGEEDN<br>MAIIKEFMRFKVHMEGSVNGHEFEIEGEGEGRPYEGTQ<br>TAKLKVTKGGPLPFAWDILSPQFMYGSKAYVKHPADIP<br>DYLKLSFPEGFKWERVMNFEDGGVVTVTQDSSLQDGE<br>FIYKVKLRGTNFPSDGPVMQKKTMGWEASSERMYPED<br>GALKGEIKQRLKLKDGGHYDAEVKTTYKAKKPVQLPG | Full length human ZO3, fused with an N-terminal MBP tag and C-terminal mCherry-His6 tag |

| | | |
|---|---|---|
| | AYNVNIKLDITSHNEDYTIVEQYERAEGRHSTGGMDEL YKLEVLFQGPGSSHHHHHHSG | |
| mCherry-CGN | MGMKIEEGKLVIWINGDKGYNGLAEVGKKFEKDTGIK VTVEHPDKLEEKFPQVAATGDGPDIIFWAHDRFGGYAQ SGLLAEITPDKAFQDKLYPFTWDAVRYNGKLIAYPIAVE ALSLIYNKDLLPNPPKTWEEIPALDKELKAKGKSALMFN LQEPYFTWPLIAADGGYAFKYENGKYDIKDVGVDNAG AKAGLTFLVDLIKNKHMNADTDYSIAEAAFNKGETAM TINGPWAWSNIDTSKVNYGVTVLPTFKGQPSKPFVGVL SAGINAASPNKELAKEFLENYLLTDEGLEAVNKDKPLG AVALKSYEEELVKDPRIAATMENAQKGEIMPNIPQMSA FWYAVRTAVINAASGRQTVDEALKDAQTNSSSNNNNN NNNNNSSGRLEVLFQGPAAAMAEPRGPVDHGVQIRFIT EPVSGAEMGTLRRGGRRPAKDARASTYGVAVRVQGIA GQPFVVLNSGEKGGDSFGVQIKGANDQGASGALSSDLE LPENPYSQVKGFPAPSQSSTSDEEPGAYWNGKLLRSHSQ ASLAGPGPVDPSNRSNSMLELAPKVASPGSTIDTAPLSS VDSLINKFDSQLGGQARGRTGRRTRMLPPEQRKRSKSL DSRLPRDTFEERERQSTNHWTSSTKYDNHVGTSKQPAQ SQNLSPLSGFSRSRQTQDWVLQSFEEPRRSAQDPTMLQF KSTPDLLRDQQEAAPPGSVDHMKATIYGILREGSSESET SVRRKVSLVLEKMQPLVMVSSGSTKAVAGQGELTRKV EELQRKLDEEVKKRQKLEPSQVGLERQLEEKTEECSRL QELLERRKGEAQQSNKELQNMKRLLDQGEDLRHGLET QVMELQNKLKHVQGPEPAKEVLLKDLLETRELLEEVLE GKQRVEEQLRLRERELTALKGALKEEVASRDQEVEHVR QQYQRDTEQLRRSMQDATQDHAVLEAERQKMSALVR GLQRELEETSEETGHWQSMFQKNKEDLRATKQELLQLR MEKEEMEEELGEKIEVLQRELEQARASAGDTRQVEVLK KELLRTQEELKELQAERQSQEVAGRHRDRELEKQLAVL RVEADRGRELEEQNLQLQKTLQQLRQDCEEASKAKMV AEAEATVLGQRRAAVETTLRETQEENDEFRRRILGLEQ QLKETRGLVDGGEAVEARLRDKLQRLEAEKQQLEEAL NASQEEEGSLAAAKRALEARLEEAQRGLARLGQEQQTL NRALEEEGKQREVLRRGKAELEEQKRLLDRTVDRLNKE | Full length human Cingulin (CGN), fused with an N-terminal MBP tag and C-terminal mCherry-His6 tag |

| | | |
|---|---|---|
| | LEKIGEDSKQALQQLQAQLEDYKEKARREVADAQRQA KDWASEAEKTSGGLSRLQDEIQRLRQALQASQAERDTA RLDKELLAQRLQGLEQEAENKKRSQDDRARQLKGLEE KVSRLETELDEEKNTVELLTDRVNRGRDQVDQLRTELM QERSARQDLECDKISLERQNKDLKTRLASSEGFQKPSAS LSQLESQNQLLQERLQAEEREKTVLQSTNRKLERKVKE LSIQIEDERQHVNDQKDQLSLRVKALKRQVDEAEEEIER LDGLRKKAQREVEEQHEVNEQLQARIKSLEKDSWRKA SRSAAESALKNEGLSSDEEFDSVYDPSSIASLLTESNLQT SSCGAPGSAGSAAGSGMVSKGEEDNMAIIKEFMRFKVH MEGSVNGHEFEIEGEGEGRPYEGTQTAKLKVTKGGPLP FAWDILSPQFMYGSKAYVKHPADIPDYLKLSFPEGFKW ERVMNFEDGGVVTVTQDSSLQDGEFIYKVKLRGTNFPS DGPVMQKKTMGWEASSERMYPEDGALKGEIKQRLKLK DGGHYDAEVKTTYKAKKPVQLPGAYNVNIKLDITSHNE DYTIVEQYERAEGRHSTGGMDELYKLEVLFQGPGSSHH HHHHSG | |
| mCherry-AFDN | MGMKIEEGKLVIWINGDKGYNGLAEVGKKFEKDTGIK VTVEHPDKLEEKFPQVAATGDGPDIIFWAHDRFGGYAQ SGLLAEITPDKAFQDKLYPFTWDAVRYNGKLIAYPIAVE ALSLIYNKDLLPNPPKTWEEIPALDKELKAKGKSALMFN LQEPYFTWPLIAADGGYAFKYENGKYDIKDVGVDNAG AKAGLTFLVDLIKNKHMNADTDYSIAEAAFNKGETAM TINGPWAWSNIDTSKVNYGVTVLPTFKGQPSKPFVGVL SAGINAASPNKELAKEFLENYLLTDEGLEAVNKDKPLG AVALKSYEEELVKDPRIAATMENAQKGEIMPNIPQMSA FWYAVRTAVINAASGRQTVDEALKDAQTNSSSNNNNN NNNNNSSGRLEVLFQGPAAAMSAGGRDEERRKLADIIH HWNANRLDLFEISQPTEDLEFHGVMRFYFQDKAAGNFA TKCIRVSSTATTQDVIETLAEKFRPDMRMLSSPKYSLYE VHVSGERRLDIDEKPLVVQLNWNKDDREGRFVLKNEN DAIPPKKAQSNGPEKQEKEGVIQNFKRTLSKKEKKEKK KREKEALRQASDKDDRPFQGEDVENSRLAAEVYKDMP ETSFTRTISNPEVVMKRRRQQKLEKRMQEFRSSDGRPDS GGTLRIYADSLKPNIPYKTILLSTTDPADFAVAEALEKY GLEKENPKDYCIARVMLPPGAQHSDEKGAKEIILDDDE CPLQIFREWPSDKGILVFQLKRRPPDHIPKKTKKHLEGK TPKGKERADGSGYGSTLPPEKLPYLVELSPGRRNHFAY YNYHTYEDGSDSRDKPKLYRLQLSVTEVGTEKLDDNSI | Full length human afadin (AFDN), fused with an N-terminal MBP tag and C-terminal mCherry-His6 tag |

| | |
|---|---|
| | QLFGPGIQPHHCDLTNMDGVVTVTPRSMDAETYVEGQ<br>RISETTMLQSGMKVQFGASHVFKFVDPSQDHALAKRSV<br>DGGLMVKGPRHKPGIVQETTFDLGGDIHSGTALPTSKST<br>TRLDSDRVSSASSTAERGMVKPMIRVEQQPDYRRQESR<br>TQDASGPELILPASIEFRESSEDSFLSAIINYTNSSTVHFKL<br>SPTYVLYMACRYVLSNQYRPDISPTERTHKVIAVVNKM<br>VSMMEGVIQKQKNIAGALAFWMANASELLNFIKQDRD<br>LSRITLDAQDVLAHLVQMAFKYLVHCLQSELNNYMPA<br>FLDDPEENSLQRPKIDDVLHTLTGAMSLLRRCRVNAAL<br>TIQLFSQLFHFINMWLFNRLVTDPDSGLCSHYWGAIIRQ<br>QLGHIEAWAEKQGLELAADCHLSRIVQATTLLTMDKY<br>APDDIPNINSTCFKLNSLQLQALLQNYHCAPDEPFIPTDL<br>IENVVTVAENTADELARSDGREVQLEEDPDLQLPFLLPE<br>DGYSCDVVRNIPNGLQEFLDPLCQRGFCRLIPHTRSPGT<br>WTIYFEGADYESHLLRENTELAQPLRKEPEIITVTLKKQ<br>NGMGLSIVAAKGAGQDKLGIYVKSVVKGGAADVDGRL<br>AAGDQLLSVDGRSLVGLSQERAAELMTRTSSVVTLEVA<br>KQGAIYHGLATLLNQPSPMMQRISDRRGSGKPRPKSEG<br>FELYNNSTQNGSPESPQLPWAEYSEPKKLPGDDRLMKN<br>RADHRSSPNVANQPPSPGGKSAYASGTTAKITSVSTGNL<br>CTEEQTPPPRPEAYPIPTQTYTREYFTFPASKSQDRMAPP<br>QNQWPNYEEKPHMHTDSNHSSIAIQRVTRSQEELREDK<br>AYQLERHRIEAAMDRKSDSDMWINQSSSLDSSTSSQEH<br>LNHSSKSVTPASTLTKSGPGRWKTPAAIPATPVAVSQPI<br>RTDLPPPPPPPPVHYAGDFDGMSMDLPLPPPPSANQIGLP<br>SAQVAAAERRKREEHQRWYEKEKARLEEERERKRREQ<br>ERKLGQMRTQSLNPAPFSPLTAQQMKPEKPSTLQRPQE<br>TVIRELQPQQQPRTIERRDLQYITVSKEELSSGDSLSPDP<br>WKRDAKEKLEKQQQMHIVDMLSKEIQELQSKPDRSAE<br>ESDRLRKLMLEWQFQKRLQESKQKDEDDEEEEDDDVD<br>TMLIMQRLEAERRARLQDEERRRQQQLEEMRKREAED<br>RARQEEERRRQEEERTKRDAEEKRRQEEGYYSRLEAER<br>RRQHDEAARRLLEPEAPGLCRPPLPRDYEPPSPSPAPGA<br>PPPPPQRNASYLKTQVLSPDSLFTAKFVAYNEEEEEEDC<br>SLAGPNSYPGSTGAAVGAHDACRDAKEKRSKSQDADS<br>PGSSGAPENLTFKERQRLFSQGQDVSNKVKASRKLTELE<br>NELNTKGAPGSAGSAAGSGMVSKGEEDNMAIIKEFMRF<br>KVHMEGSVNGHEFEIEGEGEGRPYEGTQTAKLKVTKG<br>GPLPFAWDILSPQFMYGSKAYVKHPADIPDYLKLSFPEG<br>FKWERVMNFEDGGVVTVTQDSSLQDGEFIYKVKLRGT |

| | | |
|---|---|---|
| | NFPSDGPVMQKKTMGWEASSERMYPEDGALKGEIKQR LKLKDGGHYDAEVKTTYKAKKPVQLPGAYNVNIKLDI TSHNEDYTIVEQYERAEGRHSTGGMDELYKLEVLFQGP GSSHHHHHHSG | |
| mCherry-PAR3 | MGMKIEEGKLVIWINGDKGYNGLAEVGKKFEKDTGIK VTVEHPDKLEEKFPQVAATGDGPDIIFWAHDRFGGYAQ SGLLAEITPDKAFQDKLYPFTWDAVRYNGKLIAYPIAVE ALSLIYNKDLLPNPPKTWEEIPALDKELKAKGKSALMFN LQEPYFTWPLIAADGGYAFKYENGKYDIKDVGVDNAG AKAGLTFLVDLIKNKHMNADTDYSIAEAAFNKGETAM TINGPWAWSNIDTSKVNYGVTVLPTFKGQPSKPFVGVL SAGINAASPNKELAKEFLENYLLTDEGLEAVNKDKPLG AVALKSYEEELVKDPRIAATMENAQKGEIMPNIPQMSA FWYAVRTAVINAASGRQTVDEALKDAQTNSSSNNNNN NNNNNSSGRLEVLFQGPAAAMKVTVCFGRTRVVVPCG DGHMKVFSLIQQAVTRYRKAIAKDPNYWIQVHRLEHG DGGILDLDDILCDVADDKDRLVAVFDEQDPHHGGDGTS ASSTGTQSPEIFGSELGTNNVSAFQPYQATSEIEVTPSVL RANMPLHVRRSSDPALIGLSTSVSDSNFSSEEPSRKNPTR WSTTAGFLKQNTAGSPKTCDRKKDENYRSLPRDTSNW SNQFQRDNARSSLSASHPMVGKWLEKQEQDEDGTEED NSRVEPVGHADTGLEHIPNFSLDDMVKLVEVPNDGGPL GIHVVPFSARGGRTLGLLVKRLEKGGKAEHENLFREND CIVRINDGDLRNRRFEQAQHMFRQAMRTPIIWFHVVPA ANKEQYEQLSQSEKNNYYSSRFSPDSQYIDNRSVNSAG LHTVQRAPRLNHPPEQIDSHSRLPHSAHPSGKPPSAPAS APQNVFSTTVSSGYNTKKIGKRLNIQLKKGTEGLGFSITS RDVTIGGSAPIYVKNILPRGAAIQDGRLKAGDRLIEVNG VDLVGKSQEEVVSLLRSTKMEGTVSLLVFRQEDAFHPR ELNAEPSQMQIPKETKAEDEDIVLTPDGTREFLTFEVPLN DSGSAGLGVSVKGNRSKENHADLGIFVKSIINGGAASK DGRLRVNDQLIAVNGESLLGKTNQDAMETLRRSMSTE GNKRGMIQLIVARRISKCNELKSPGSPPGPELPIETALDD RERRISHSLYSGIEGLDESPSRNAALSRIMGESGKYQLSP TVNMPQDDTVIIEDDRLPVLPPHLSDQSSSSHDDVGFV TADAGTWAKAAISDSADCSLSPDVDPVLAFQREGFGRQ SMSEKRTKQFSDASQLDFVKTRKSKSMDLGIADETKLN | Full length human PAR3, fused with an N-terminal MBP tag and C-terminal mCherry-His6 tag |

| | | |
|---|---|---|
| | TVDDQKAGSPSRDVGPSLGLKKSSSLESLQTAVAEVTL NGDIPFHRPRPRIIRGRGCNESFRAAIDKSYDKPAVDDD DEGMETLEEDTEESSRSGRESVSTASDQPSHSLERQMNG NQEKGDKTDRKKDKTGKEKKKDRDKEKDKMKAKKG MLKGLGDMFRFGKHRKDDKIEKTGKIKIQESFTSEEERI RMKQEQERIQAKTREFRERQARERDYAEIQDFHRTFGC DDELMYGGVSSYEGSMALNARPQSPREGHMMDALYA QVKKPRNSKPSPVDSNRSTPSNHDRIQRLRQEFQQAKQ DEDVEDRRRTYSFEQPWPNARPATQSGRHSVSVEVQM QRQRQEERESSQQAQRQYSSLPRQSRKNASSVSQDSWE QNYSPGEGFQSAKENPRYSSYQGSRNGYLGGHGFNAR VMLETQELLRQEQRRKEQQMKKQPPSEGPSNYDSYKK VQDPSYAPPKGPFRQDVPPSPSQVARLNRLQTPEKGRPF YSGAPGSAGSAAGSGMVSKGEEDNMAIIKEFMRFKVH MEGSVNGHEFEIEGEGEGRPYEGTQTAKLKVTKGGPLP FAWDILSPQFMYGSKAYVKHPADIPDYLKLSFPEGFKW ERVMNFEDGGVVTVTQDSSLQDGEFIYKVKLRGTNFPS DGPVMQKKTMGWEASSERMYPEDGALKGEIKQRLKLK DGGHYDAEVKTTYKAKKPVQLPGAYNVNIKLDITSHNE DYTIVEQYERAEGRHSTGGMDELYKLEVLFQGPGSSHH HHHHSG | |
| mCherry-CDH1-IC | MGSSHHHHHHSSGRLEVLFQGPMVSKGEEDNMAIIKEF MRFKVHMEGSVNGHEFEIEGEGEGRPYEGTQTAKLKVT KGGPLPFAWDILSPQFMYGSKAYVKHPADIPDYLKLSFP EGFKWERVMNFEDGGVVTVTQDSSLQDGEFIYKVKLR GTNFPSDGPVMQKKTMGWEASSERMYPEDGALKGEIK QRLKLKDGGHYDAEVKTTYKAKKPVQLPGAYNVNIKL DITSHNEDYTIVEQYERAEGRHSTGGMDELYKGSAGSA AGSGAAAVRRRRVVKEPLLPPEDDTRDNVYYYDEEGG GEEDQDFDLSQLHRGLDARPEVTRNDVAPTLLSVPQYR PRPANPDEIGNFIDENLKAADTDPTAPPYDSLLVFDYEG SGSEAASLSSLNSSESDQDQDYDYLNEWGNRFKKLAD MYGGGEDD | Human cytosolic fragments (734-885), fused with an N-terminal His6-mCherry tag |
| JAM-A-IC | MGSSHHHHHHHHHHSSGRLEVLFQGPAAAYRRGYFDR AKKGTSSKKVIYSQPAARSEGEFRQTSSFLV | Dog cytosolic fragments (260-298), fused with an |

| | | |
|---|---|---|
| | | N-terminal His10-tag |
| mCherry-CRB3-IC | MGSSHHHHHHSSGRLEVLFQGPMVSKGEEDNMAIIKEF MRFKVHMEGSVNGHEFEIEGEGEGRPYEGTQTAKLKVT KGGPLPFAWDILSPQFMYGSKAYVKHPADIPDYLKLSFP EGFKWERVMNFEDGGVVTVTQDSSLQDGEFIYKVKLR GTNFPSDGPVMQKKTMGWEASSERMYPEDGALKGEIK QRLKLKDGGHYDAEVKTTYKAKKPVQLPGAYNVNIKL DITSHNEDYTIVEQYERAEGRHSTGGMDELYKGSAGSA AGSGAAARKLREKRQTEGTYRPSSEEQVGARVPPTPNL KLPPEERLI | Human cytosolic fragments (91-120), fused with an N-terminal His6-mCherry tag |
| SNAP-Utrophin | MDKDCEMKRTTLDSPLGKLELSGCEQGLHEIKLLGKGT SAADAVEVPAPAAVLGGPEPLMQATAWLNAYFHQPEA IEEFPVPALHHPVFQQESFTRQVLWKLLKVVKFGEVISY QQLAALAGNPAATAAVKTALSGNPVPILIPCHRVVSSSG AVGGYEGGLAVKEWLLAHEGHRLGKPGLGPAGIGAPG SMAKYGEHEASPDNGQNEFSDIIKSRSDEHNDVQKKTF TKWINARFSKSGKPPINDMFTDLKDGRKLLDLLEGLTG TSLPKERGSTRVHALNNVNRVLQVLHQNNVELVNIGGT DIVDGNHKLTLGLLWSIILHWQVKDVMKDVMSDLQQT NSEKILLSWVRQTTRPYSQVNVLNFTTSWTDGLAFNAV LHRHKPDLFSWDKVVKMSPIERLEHAFSKAQTYLGIEK LLDPEDVAVRLPDKKSIIMYLTSLFEVLPQQVTID | N-terminal part of human Utrophin (1-261), fused with a C-terminal SNAP tag |

**Supplement movies:**

**Movie S1. ZO1 surface Condensation.** 1400 molecules/ $\mu m^2$ 14-mer receptors (labeled with 5% DyLight 650, Magenta) were anchored to DGS-NTA(Ni) containing lipid membranes. Images were taken every 10s for 15 min after adding 200 nM ZO1-mGFP (Green). ZO1 was gradually recruited to the membrane, and de-mixed to form a dense phase and a dilute phase. During this process, Ostwald ripening and fusion events happened to form surface condensates.

**Movie S2. Local actin polymerization.** 450 molecules/ $\mu m^2$ 14-mer receptor (labeled with DyLight 550, Magenta) was anchored to DGS-NTA(Ni) containing lipid membrane. 200 nM ZO1-GFP (Green) was added for 10 min to form ZO1 surface condensates. 3 µM G-actin (labeled with 5% Alexa Fluor 647, Yellow) was added to ZO1 surface condensates. Images were taken every 30 s for 60 min. G-actin was recruited to ZO1 surface condensates, polymerized out and bundled. Actin bundles connected with each other to form a network. Simultaneously, ZO1 surface condensates spread on actin bundles to form a continuous receptor-ZO1-actin network.

**Movie S3. Local actin bundling.** Actin bundling during the time was quantified from movie S2. Actin polymerized from the ZO1 surface condensates as a single filament. New actin filaments grew on top of the F-actin filaments to form actin bundles.

**References:**

48. H. Sun, Y. Luo, Y. Miao, Purification of Globular Actin from Rabbit Muscle and Pyrene Fluorescent Assays to Investigate Actin Dynamics in vitro. *Bio Protoc* **8**, e3102 (2018).
49. M. Swoboda *et al.*, Enzymatic Oxygen Scavenging for Photostability without pH Drop in Single-Molecule Experiments. *Acs Nano* **6**, 6364-6369 (2012).

### Supplementary Information on Theoretical Model:
### Assembly of tight junction belts by surface condensation and actin elongation

#### CONTENTS

A. Model for protein binding to receptors in a membrane 2

B. Thermodynamics and molecular interactions 3

C. Equilibrium thermodynamics 5

D. Binding fraction 6

E. Non-equilibrium thermodynamics with membrane binding 7

F. Parameter choices and parameters obtained from fitting to the experimental data 8

G. Derivation of dilute binding affinity given in Eq. (23) 14

H. Kinetics of membrane phase separation 14

References 15

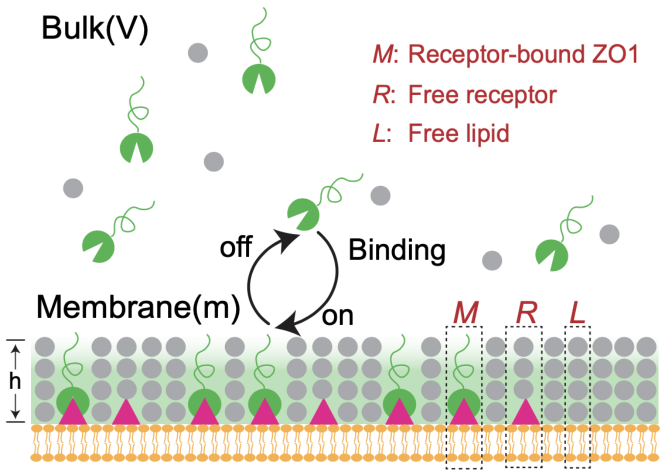

FIG. 1: **Schematics of the bulk-membrane coupled system.** The receptor (red) anchors to the lipid (orange) of supported lipid bilayers (SLBs). Proteins (green) can bind to receptors, forming a receptor-protein complex $M$. We introduce two more effective components in the membrane: $R$ denotes lipids with a receptor without any protein bound but a water layer on top, and $L$ represents a lipid patch without a receptor but with the water layer on top. The effective components $M$, $R$, and $L$ are restricted to the membrane.

#### A. Model for protein binding to receptors in a membrane

In our model, we consider three spatial domains (see Fig. 1): the bulk containing proteins and solvent, and the membrane composed of lipids and receptors with a layer of proteins bound to the receptors in the membrane. The proteins in bulk can bind to receptors that anchor lipids in the membrane, while free lipids lacking a receptor are not accessible for protein binding. Since receptors are confined to the membrane, bound proteins can only move within a layer adjacent to the membrane surface. This layer also contains solvent molecules, which are exchanged with the bulk when proteins bind. Note that the bulk does not contain any receptor.

To describe the setting outlined above, we propose a spatially coarse-grained model for the membrane and the adjacent layer. To this end, we define three effective components on a surface: lipid patches with receptors bound to protein $M$, lipid patches with receptors but without protein-bound $R$, and lipid patches with only the water layer on top $L$. As illustrated in Fig. 1, we consider that an effective component $M$ is composed of $q$ receptors, and one protein $B$. We also introduce the molecular volume ratio $n_b$ between proteins $B$ and solvent $S$. The binding for the coarse-grained model can be written as

$$B + q\,R \rightleftharpoons M + n_b\,S\,. \tag{3}$$

Conservation of volume implies for the molecules' molecular volumes,

$$\nu_b + q\,\nu_r\,h = \nu_m\,h + n_b\,\nu_s\,, \tag{4}$$

where $h$ is the height of coarse-grained surface (see Fig. 1). $\nu_m$, $\nu_r$, $\nu_l$ are the area fractions of the membrane components $M$, $R$, $L$, respectively and $\nu_b$ is the volume fraction of protein in the bulk, $B$. For incompressibility in the bulk, we set $\nu_b = n_b\,\nu_s$, and for incompressibility in the layer adjacent the membrane, $n_m = q\,n_r$, where $n_r = \nu_r/\nu_l$, $n_m = \nu_m/\nu_l$. In our study, scaffold proteins are considered to have a larger area fraction than receptors. In the coarse-grained surface, we set $\nu_m = \nu_r$. Note that the actual area fraction of protein equals $m\,\nu_m$, where $m$ is the ratio between real area fraction of scaffold protein and $\nu_r$.

#### B. Thermodynamics and molecular interactions

The thermodynamics of the system is characterized by bulk volume fraction $\phi(\boldsymbol{x}, t)$ of the proteins, volume fraction $\phi_s(\boldsymbol{x}, t)$ of the solvent, and the area fraction of ZO1-receptor complex that is confined in the membrane, $\phi_m(\boldsymbol{x}, t)$, the free receptor $\phi_r(\boldsymbol{x}, t)$. and the free lipid patch $\phi_l(\boldsymbol{x}, t)$. The volume fraction field depend on position $\boldsymbol{x} = (x, y, z)$. The membrane is located at $z = 0$. All area fraction fields depend on position $\boldsymbol{x} = (x, y)$. We can reduce the number of independent degrees of freedom exploiting incompressibility both in the bulk, $\phi + \phi_s = 1$, and at the membrane surface $\phi_m + \phi_r + \phi_l = 1$, respectively.

The total free energy $F$ of the system is composed of different contributions: the free energy density $f_b(\phi)$ in the bulk $V$, the free energy density $f_m(\phi_m, \phi_r)$ at the membrane surface $m$, and coupling between bulk and membrane $J(\phi, \phi_m, \phi_r)$. The total free energy reads:

$$\begin{aligned} F[\phi, \phi_m, \phi_r] = \int_V d^3x \Big[f_b(\phi) + \frac{1}{2}\kappa|\nabla\phi|^2\Big] + \int_m d^2x \Big[f_m(\phi_m, \phi_r) + J(\phi, \phi_m, \phi_r) \\ + \frac{1}{2}\kappa_r|\nabla_{||}\phi_r|^2 + \frac{1}{2}\kappa_m|\nabla_{||}\phi_m|^2 + \kappa_{mr}\nabla_{||}\phi_r \cdot \nabla_{||}\phi_m\Big]\,. \end{aligned} \tag{5}$$

The free energy penalties due to concentration gradients is characterized by $\kappa$, $\kappa_m$, $\kappa_r$, and $\kappa_{mr}$ in bulk and membrane, respectively, and $\nabla_{||} = (\partial_x, \partial_y)$ denotes the gradient parallel to the membrane plane.

The coupling free energy between the bulk and the membrane is written as:

$$J(\phi, \phi_m, \phi_r) = \frac{k_B T}{\nu_l} \Big[ \chi_{bl}\, \phi\, (1 - \phi_r - \phi_m) + \chi_{br}\, \phi\, \phi_r + \chi_{bm}\, \phi\, \phi_m \Big] , \tag{6}$$

where the interactions of proteins in the bulk with free lipid patch, with free receptors and with membrane-bound proteins at the membrane are described by $\chi_{bl}$, $\chi_{br}$ and $\chi_{bm}$, respectively. We use Flory-Huggins free energies in the bulk and membrane, respectively:

$$f_b(\phi) = \frac{k_B T}{\nu_s} \Big[ \frac{\phi}{n_b} \ln \phi + (1 - \phi) \ln (1 - \phi) + \chi_b \phi (1 - \phi) + \omega_b \phi \Big] , \tag{7}$$

$$f_m(\phi_m, \phi_r) = \frac{k_B T}{\nu_l} \Big[ \frac{\phi_m}{n_m} \ln \phi_m + \frac{\phi_r}{n_r} \ln \phi_r + (1 - \phi_m - \phi_r) \ln(1 - \phi_m - \phi_r) \tag{8}$$

$$+ \chi_{mr} \phi_m\, \phi_r + \chi_{ml} \phi_m (1 - \phi_m - \phi_r) + \chi_{rl} \phi_r (1 - \phi_m - \phi_r) + \omega_m \phi_m + \omega_r \phi_r \Big] .$$

We take the internal free energy of proteins in the bulk as $\omega_b$, denote the internal free energies of receptor-protein complex and free receptor as $\omega_m$ and $\omega_r$. We note that $\nu_l$ represents the area of an effective free lipid patch to avoid considering molecules of very different size. As a consequence, to obtain the actual Flory-Huggins parameters corresponding to the lipid-ZO1 interaction, it need to be rescaled by the number of lipids in each lipid patch.

If the free lipid area fraction is zero ($\phi_l = 0$), our model reduces to the binary mixture in the model discussed recently [5]. Without receptor in the membrane ($\phi_r + \phi_m = 0$), our model yield two decoupled systems: a two dimensional membrane and a three dimensional binary mixture in the bulk. If $\phi_l > 0$ and $\phi_r > 0$, our model corresponds to a ternary mixture in the membrane coupled to a binary mixture in the bulk. This model accounts for one more component, i.e., the receptor-protein complex in comparison to the thermodynamic model developed in the paper [3]. In addition, our model describes the binding/unbinding events of proteins to the free receptors (or tethers) anchored in the membrane.

Due to incompressibility of bulk and membrane, we obtain exchange chemical potentials of the proteins in the bulk $\mu$, the receptor-protein complex $\mu_m$ and the free receptor in the membrane $\mu_r$:

$$\mu \equiv \nu_b \frac{\delta F}{\delta \phi} = \nu_b \left( \frac{\partial f}{\partial \phi} - \kappa \nabla^2 \phi \right) , \tag{9a}$$

$$\mu_m \equiv \nu_m \frac{\delta F}{\delta \phi_m} = \nu_m \left( \frac{\partial f_m}{\partial \phi_m} + \frac{\partial J}{\partial \phi_m} - \kappa_m \nabla_{||}^2 \phi_m - \kappa_{mr} \nabla_{||}^2 \phi_r \right) , \tag{9b}$$

$$\mu_r \equiv \nu_r \frac{\delta F}{\delta \phi_r} = \nu_r \left( \frac{\partial f_m}{\partial \phi_r} + \frac{\partial J}{\partial \phi_r} - \kappa_r \nabla_{||}^2 \phi_r - \kappa_{mr} \nabla_{||}^2 \phi_m \right) . \tag{9c}$$

We consider a finite system where the bulk has a volume $V = L_z L^2$, and $L_z$ denotes the height orthogonal to the membrane surface which has surface areas of $L^2$. For simplicity, we set $L_z = L$ in this study.

#### C. Equilibrium thermodynamics

In this section, we derive the conditions at thermodynamic equilibrium. At thermodynamic equilibrium, the total free energy $F$ is minimal. There are constraints for the minimization since total number of proteins $N$ and the total number of receptors at the membrane $N_r$ are conserved. The minimum of the free energy fulfills

$$0 = \delta \left\{ F - \lambda \Big[ \int_V d^3x \, \phi/\nu_b + \int_m d^2x \, \phi_m/\nu_m - N \Big] - \lambda_r \Big[ \int_m d^2x \, \Big( \phi_r/\nu_r + q \, \phi_m/\nu_m \Big) - N_r \Big] \right\} , \tag{10}$$

where $\lambda$ and $\lambda_r$ denote Lagrange multipliers corresponding to the two respective conservation laws. Using Eq. (5), the variation of the total free energy is given as

$$\begin{aligned}
\delta F = & \int_V d^3x \Big( \frac{\partial f}{\partial \phi} - \kappa \nabla^2 \phi \Big) \delta\phi \\
& + \int_m d^2x \Big( \frac{\partial f_m(\phi_m, \phi_r)}{\partial \phi_m} - \kappa_m \nabla_{||}^2 \phi_m - \kappa_{mr} \nabla_{||}^2 \phi_r + \frac{\partial J}{\partial \phi_m} \Big) \delta\phi_m \\
& + \int_m d^2x \Big( \frac{\partial f_m(\phi_m, \phi_r)}{\partial \phi_r} - \kappa_r \nabla_{||}^2 \phi_r - \kappa_{mr} \nabla_{||}^2 \phi_m + \frac{\partial J}{\partial \phi_r} \Big) \delta\phi_r \\
& + \int_m \Big( \frac{\partial J}{\partial \phi} \Big) \delta\phi + \int_{m,\partial V} \Big( \kappa \, \boldsymbol{n} \cdot \nabla \phi \Big) \delta\phi \\
& + \int_{\partial m} dx \, \boldsymbol{t} \cdot \Big( \kappa_m \nabla_{||} \phi_m + \kappa_{mr} \nabla_{||} \phi_r \Big) \delta\phi_m + \int_{\partial m} dx \, \boldsymbol{t} \cdot \Big( \kappa_r \nabla_{||} \phi_r + \kappa_{mr} \nabla_{||} \phi_m \Big) \delta\phi_r \, ,
\end{aligned} \tag{11}$$

where $V$ is the bulk volume, membrane surface $m$, and $\partial V$ its boundary except the membrane surface. $\partial m$ is the boundary contour of membrane surfaces.

Using Eqs. (9), we identify $\mu_m = \lambda + q \, \lambda_r$, $\mu = \lambda$ and $\mu_r = \lambda_r$. The entire system composed of membrane and bulk is at thermodynamic equilibrium if binding equilibrium is satisfied:

$$\mu + q \, \mu_r = \mu_m \, . \tag{12a}$$

Binding equilibrium suggest to introduce the binding energy per receptor-protein complex $M$.

$$\Delta\omega = n_m \, \omega_m - q \, n_r \, \omega_r - n_b \, \omega_b \, , \tag{12b}$$

If two phases, denoted as I and II, coexist in the membrane, we have the equilibrium conditions:

$$\mu_m^{\mathrm{I}} = \mu_m^{\mathrm{II}}, \tag{12c}$$

$$\mu_r^{\mathrm{I}} = \mu_r^{\mathrm{II}}, \tag{12d}$$

$$f^{\mathrm{I}} - f^{\mathrm{II}} = \mu_m^{\mathrm{I}} (\phi_m^{\mathrm{I}} - \phi_m^{\mathrm{II}}) + \mu_r^{\mathrm{I}} (\phi_r^{\mathrm{I}} - \phi_r^{\mathrm{II}}) \, . \tag{12e}$$

The total amount of receptor in the membrane is conserved:

$$A\phi_r^0 = A^{\mathrm{I}}(\phi_m^{\mathrm{I}} + \phi_r^{\mathrm{I}}) + (A - A^{\mathrm{I}})(\phi_m^{\mathrm{II}} + \phi_r^{\mathrm{II}}) \,, \tag{12f}$$

where $A^{\mathrm{I}}$ is the surface condensate area in the membrane, $A = L^2$ is the total area of the membrane, and $\phi_r^0$ is the initial area fraction of the receptor in the membrane.

The boundary conditions at the membrane $m$ and other boundaries enclosing the bulk, $\partial V$, at thermodynamic equilibrium are:

$$\boldsymbol{n} \cdot \nabla\phi = -\kappa^{-1}\frac{\partial J}{\partial \phi} \,, \qquad x \in m \,, \tag{13a}$$

$$\boldsymbol{n} \cdot \nabla\phi = 0 \,, \qquad x \in \partial V \,, \tag{13b}$$

$$\boldsymbol{t} \cdot \Big(\kappa_m \nabla_{||}\phi_m + \kappa_{mr}\nabla_{||}\phi_r\Big) = 0 \,, \qquad x \in \partial m \,, \tag{13c}$$

$$\boldsymbol{t} \cdot \Big(\kappa_r \nabla_{||}\phi_r + \kappa_{mr}\nabla_{||}\phi_m\Big) = 0 \,, \qquad x \in \partial m \,. \tag{13d}$$

The first condition characterizes the interactions with the membrane surface and thereby also determine wetting behavior of bulk condensates. The second conditions specifies that the remaining boundaries of the cubic system are neutral, i.e., the free energy is not affected by increasing or decreasing the bulk volume fractions adjacent to $\partial V$. The last two conditions describe the perimeters of the membrane boundary $\partial m$ as neutral.

#### D. Binding fraction

In the experiments, we observed that maximal fraction of bound receptors is different for different oligomerization states of the receptor, i.e., monomer and 14-mer. Specifically, while all 14-mer receptors can be bound by proteins from the bulk, only a fraction of monomer receptors are available for binding. This phenomena was already reported in Ref. [2] and tracked back to a molecular switch mechanism.

We quantify the binding fraction $c$ as the ratio between total membrane-bound protein area fraction and total free receptor area fraction at saturated state, i.e. $c = \bar{\phi}_m/\phi_r^0$. Based on the experimental data, we obtain that $c = 1$ for the 14-mer receptors and $c = 0.014$ for monomer receptors. In our model, we include the reduced binding fraction mechanism in the following way:

We calculate the membrane-bound protein area fraction $\phi_m$ from the thermodynamic conditions (12). Then, (1) If $\phi_m \leq c\phi_r^0$, a finite binding fraction $c$ has no effects. (2) If $\phi_m > c\phi_r^0$, we use

$$\phi_m = c\,\phi_r^0\,,$$
$$\phi_r = \phi_r^0 - c\,\phi_r^0\,,$$

where $\phi_r^0$ is the average area fraction of the free receptors in the membrane initially.

#### E. Non-equilibrium thermodynamics with membrane binding

In this section, we discuss the kinetic equations for membrane binding and membrane phase separation that govern the relaxation toward thermodynamic equilibrium. This derivations are based on irreversible thermodynamics and the details of the derivation is presented in a separate work. The kinetic equations are used to verify that ZO1-receptor rich domains can develop from the experiential initial conditions of a ZO1-free membrane surface. Moreover, the kinetics is used to compute the the spatial patterns of ZO1-receptor surface condensates (see main text, Fig. 2(C,E)). Note that the phase diagram in the main text (Fig. 2(G,H)) is calculated at equilibrium corresponding to $\partial_t\phi = 0$, $\partial_t\phi_m = 0$ and $\partial_t\phi_r = 0$, or equivalently, Eq. (12) when the system is large enough.

The governing kinetic equations for the bulk volume fraction $\phi$, the area fraction of receptor-protein complexes $\phi_m$ and free receptor in the membrane $\phi_r$ read:

$$\partial_t\phi = D_b\,\nabla\cdot(\phi(1-\phi)\nabla\mu)\,, \qquad x\in V\,, \tag{14a}$$
$$\partial_t\phi_m = D_m\,\nabla\cdot(\phi_m(1-\phi_m-\phi_r)\nabla\mu_m) - \Lambda_s\left[\exp\left(\mu + q\,\mu_r\right) - \exp\left(\mu_m\right)\right]\,, \qquad x\in m\,, \tag{14b}$$
$$\partial_t\phi_r = D_r\,\nabla\cdot(\phi_r(1-\phi_r-\phi_m)\nabla\mu_r) + q\frac{\nu_r}{\nu_m}\Lambda_s\left[\exp\left(\mu + q\,\mu_r\right) - \exp\left(\mu_m\right)\right]\,, \qquad x\in m\,, \tag{14c}$$

with the boundary equations

$$\boldsymbol{n}D_b\cdot\phi(1-\phi)\nabla\mu = \frac{\nu_b}{\nu_m}\left[\exp\left(\mu + q\,\mu_r\right) - \exp\left(\mu_m\right)\right]\,, \qquad x\in m\,, \tag{14d}$$
$$\boldsymbol{n}\cdot\kappa\nabla\phi = -\frac{\partial J}{\partial\phi}\,, \qquad x\in m\,, \tag{14e}$$
$$-\boldsymbol{n}\cdot(D_b\phi(1-\phi)\nabla\mu) = 0\,, \qquad x\in\partial V\,, \tag{14f}$$
$$\boldsymbol{n}\cdot\nabla\phi = 0\,, \qquad x\in\partial V\,, \tag{14g}$$

where $D_b$, $D_m$ and $D_r$ are the diffusion coefficients of scaffold protein in the bulk, receptor-protein complex and free receptor in the membrane. Moreover, $\Lambda_s$ is the binding rate of scaffold protein ZO1 binding/unbinding to the membrane.

To solve such equations numerically we rescale length $x \to x/\ell$, where $\ell = 10\,\mu m$ is the experimental field of view, and rescale time $t \to t \cdot \Lambda_s^{-1}$. The non-dimensional quantities are:

$$\tilde{J}(\phi, \phi_m, \phi_r) = J(\phi, \phi_m, \phi_r)\frac{\nu_l}{k_{\mathrm{B}}T}\,, \tag{15}$$

$$\tilde{\kappa}_m = \frac{1}{\ell^2}\kappa_m\frac{\nu_m}{k_{\mathrm{B}}T} = \frac{\nu_m}{\ell^2}\,, \quad \tilde{\kappa} = \frac{1}{\ell^2}\kappa\frac{\nu_b}{k_{\mathrm{B}}T} = \frac{\nu_b^{2/3}}{\ell^2}\,, \quad \bar{\kappa} = \kappa\frac{\nu_l}{k_{\mathrm{B}}T}\frac{1}{\ell} = \frac{\nu_l}{\nu_b^{1/3}\ell}\,, \tag{16}$$

where $\tilde{J}(\phi, \phi_m, \phi_r)$ is the non-dimensional binding flux between membrane and bulk. $\tilde{\kappa}_m$, $\tilde{\kappa}$, and $\bar{\kappa}$ are the non-dimensional gradient coefficients in the 2D membrane, 3D bulk and bulk boundaries adjacent to the membrane. We also introduce the non-dimensional parameters:

$$\mathcal{D}_i = \frac{D_i}{\Lambda_s \ell^2}\,, \tag{17}$$

where $i = b, m, r$. Here, $\mathcal{D}_i$ are inverse "Damköhler numbers", which compares the time-scale of a chemical reaction, i.e., binding, with the time-scale of diffusive transport.

With the non-dimensional quantities, we achieve the following non-dimensional equations:

$$\partial_t \phi = \mathcal{D}_b \nabla \cdot (\phi(1-\phi)\nabla\tilde{\mu})\,, \qquad x \in V\,, \tag{18a}$$

$$\partial_t \phi_m = \mathcal{D}_m \nabla \cdot (\phi_m(1-\phi_m-\phi_r)\nabla\tilde{\mu}_m) - [\exp(\tilde{\mu} + q\,\tilde{\mu}_r) - \exp(\tilde{\mu}_m)]\,, \qquad x \in m\,, \tag{18b}$$

$$\partial_t \phi_r = \mathcal{D}_r \nabla \cdot (\phi_r(1-\phi_r-\phi_m)\nabla\tilde{\mu}_r) + q\frac{\nu_r}{\nu_m}[\exp(\tilde{\mu} + q\,\tilde{\mu}_r) - \exp(\tilde{\mu}_m)]\,, \qquad x \in m\,, \tag{18c}$$

with the non-dimensional boundary equations

$$\boldsymbol{n}\mathcal{D}_b \cdot \phi(1-\phi)\nabla\tilde{\mu} = \frac{1}{\ell}\frac{\nu_b}{\nu_m}[\exp(\tilde{\mu} + q\,\tilde{\mu}_r) - \exp(\tilde{\mu}_m)]\,, \qquad x \in m\,, \tag{18d}$$

$$\boldsymbol{n} \cdot \bar{\kappa}\nabla\phi = -\frac{\partial\tilde{J}}{\partial\phi}\,, \qquad x \in m\,, \tag{18e}$$

$$-\boldsymbol{n} \cdot (\mathcal{D}_b\phi(1-\phi)\nabla\tilde{\mu}) = 0\,, \qquad x \in \partial V\,, \tag{18f}$$

$$\boldsymbol{n} \cdot \nabla\phi = 0\,, \qquad x \in \partial V\,. \tag{18g}$$

In the equations above, chemical potentials and coupling free energy $\tilde{J}$ are measured in units of $k_BT$, $\tilde{\mu} \equiv \mu/(k_BT)$ and $\tilde{J} \equiv \nu_m J/(k_BT)$. All the parameter values and their dimensionless values in the following study are given in Table VI.

#### F. Parameter choices and parameters obtained from fitting to the experimental data

We use our theoretical model Eqs. (14) to understand the mechanism for membrane phase separation. According to the experimental studies, membrane phase separation occurs for bulk

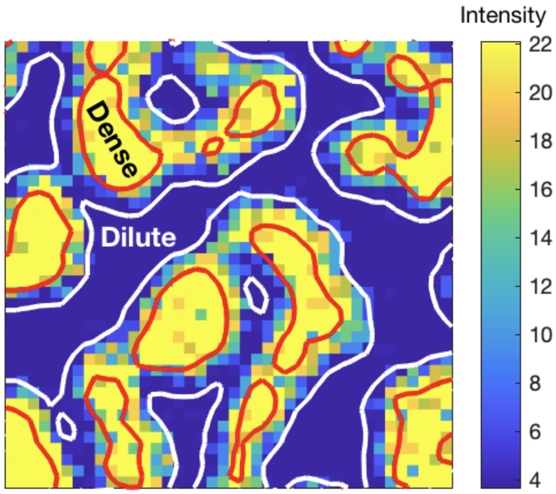

FIG. 2: **Illustration of the mask for receptor intensity to determine the surface concentrations inside and outside of the surface condensates in the membrane.** The blue area represents the dilute phase and the yellow area the dense phase.

protein concentrations of a few tens of $nM$. For concentrations in the order of 10 $nM$, there is no phase separation in the bulk since the bulk saturation concentration is about $10\,\mu M$. Thus, we simplify our model by adapting it in an approximate fashion to these experimental conditions by setting bulk interaction parameter $\chi_b = 0$ and use, for simplicity, equivalent solvent and solute molecule size, i.e., $n_b = 1$. In particular, we choose the coupling coefficients in Eq. (6) as $\chi_{bl} = \chi_{bm} = \chi_{br} = 0$, implying that the profile along $z$, $\phi(z)$ is constant at thermodynamic equilibrium. Note that during the time evolution, the bulk volume fraction described by Eq. (18a) can be heterogeneous. At thermodynamic equilibrium, however, the bulk can be thought as a finite homogeneous reservoir for scaffold proteins.

We roughly estimated the size of 14-mer receptor and monomer receptor from the structure in Pymol based on their structures [1, 4].

$$d_{\text{14-mer Receptor}} = 15\,nm\,, \tag{19}$$

$$d_{\text{monomer Receptor}} = 5\,nm\,, \tag{20}$$

The corresponding molecule diameter of the ZO1 protein is achieved from AFM (see Fig. 1I in main text),

$$d_{\text{ZO1}} = 15\,nm\,, \tag{21}$$

We choose the size of lipid patch in the membrane as:

$$d_{\text{lipid}} = 5\,nm\,. \tag{22}$$

This is an effective free lipid patch to avoid considering molecules of very different size as we mentioned before. We note that the molecule area ratio for 14-mer receptor are given $n_m = n_r = 9$ with lipid patch as a reference component in the membrane, while the ones for monomer are $n_m = n_r = 1$.

With the molecule size given above, we can estimate the volume fraction of the ZO1 proteins in the bulk. We find that the bulk volume fraction value is very tiny (roughly $10^{-5}$). From a numerical perspective, such low numerical volume fractions requires a very fine discretization in both time and space to sample the curvature of the free energy density. The large curvature in volume fraction arises from the entropic term and mathematically speaking, from the logarithmic dependence in the free energy. To avoid extremely fine numerical grids and thus unnecessary computational resources, we scale the bulk volume fraction $\phi^{\text{raw}}$ with a scaling factor such that the volume fraction $\phi$ equals to 0.1 when the ZO1-protein number density is $50\,nM$ in the bulk: $\phi = C_0\,\phi^{\text{raw}}$, where we use $C_0 = 8146$.

Based on the discussions above, there are two unknown parameters in our model: the interaction parameter between membrane-bound proteins and free lipid, $\chi_{ml}$, and the internal free energy difference between the bound and unbound state, $\omega_m$. We can equivalently consider the following parameter pair: $(\chi_{ml},\,\Omega)$, where

$$\Omega = -\left(n_m\,\chi_{ml}\left(1-\phi_r^0\right) + n_m\,\omega_m\right)\,, \tag{23}$$

is the dilute binding affinity; $\phi_r^0$ is the average receptor area fraction. Please note that in the main text and the following study, we abbreviate $\chi_{ml}$ as $\chi$, and $\omega_m$ as $\omega$; Eq. (23) is obtained in the dilute limit of our model, where there are no phase transitions and solely binding between bulk and membrane surface binding; derivation see Section G.

To obtain the two remaining model parameters $(\chi,\,\omega)$ (and equivalently, $(\chi,\,\Omega)$), we fit our thermodynamic model (Eq. (12)) to the experimental data for the receptor-protein area fraction $\phi_m$ in the membrane. To this end, we analyse the microscope images and compute the density of the components in the membrane and bulk. For this purpose, we define a mask to find the dense and dilute phases, respectively. In specific, we use a Gaussian filter with a two-dimensional Gaussian smoothing kernel with standard deviation of 0.8 to smooth the raw receptor intensity data. Then, we set the area with $I_{\text{receptor}} \geq 22$ for dense phase, while $I_{\text{receptor}} \leq 4$ for dilute phase. The area in between is the interface area (see Fig. 2).

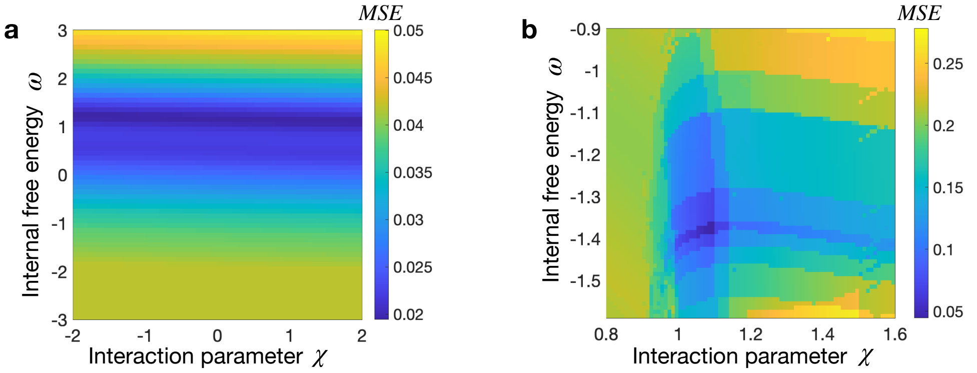

FIG. 3: **Mean squared error ($MSE$) color maps with respect to interaction parameters $\chi$ and internal free energy $\omega$ for monomer receptor (a) and 14-mer receptor (b).** For the considered parameter range, $MSE$ is minimized for 14-mer at $\chi = 1.09$ and $\omega = -1.37$, and for monomer at $\chi = -0.4$ and $\omega = 1.1$. We note that though there is a minimum of the $MSE$ corresponding to at $\chi = -0.4$, the color map in figure (a) suggests that the $\chi$ value for the monomer cannot be determined as there is a "valley" of similar $MSE$ along $\chi$. Detailed discussion see below.

For fitting, we consider two sets of experiment data, one for varying bulk ZO1 concentration and another for varying receptor amount in the membrane. We solve the optimal values of the interaction parameter $\chi$ and internal free energy coefficient $\omega$ by minimizing the mean squared error:

$$MSE = \frac{1}{N} \sum_{i=1}^{N} |\phi_{\text{ZO1}}^{\text{data}} - \phi_{\text{ZO1}}^{\text{simu}}| \,, \tag{24}$$

where $N$ is the number of total data points in the experiments. Moreover, $\phi_{\text{ZO1}}^{\text{data}}$ is the area fraction of membrane-bound ZO1. We solve $\phi_{\text{ZO1}}^{\text{simu}}$ from the equilibrium condition Eq. (12). For a range in $\chi$ and $\omega$ (see Fig. 3), we obtain the minimized $MSE$ for 14-mer with values $\chi = 1.09, \omega = -1.37$. We compare the simulated results with experimental data (see Fig. 2B and Fig. 2D in main text). Good agreement is observed in term of equilibrium densities and transition lines. Similarly, we obtain the values $\chi = -0.4, \omega = 1.1$ for monomer. It should be acknowledged that despite the existence of a $MSE$ minimum at an interaction parameter $\chi = -0.4$, the color map of $MSE$ depicted in Fig. 3(a) shows a "valley" of closely-matched $MSE$ values along the considered $\chi$ range. This phenomenon can be attributed to the fact that the interaction parameter $\chi$ hardly

| **Parameter name** | **Symbol** | **Value [units]** | **Value** |
|---|---|---|---|
| **Characteristic scales** | | | |
| Time scale | $t_0 = \Lambda_s^{-1}$ | 100 $[s]$ | 1 |
| Length scale | $l$ | 10 $[\mu m]$ | 1 |
| **Receptor valency** | | | |
| Number of receptors per ZO1 bound | $q$ | $\nu_m/\nu_r$ | 1 |
| **Molecular area/volume** | | | |
| Molecular area of free lipid in the membrane | $\nu_l$ | $2.5^2\pi[nm^2]$ | $2.5^2\pi/l^2$ |
| Molecular area of bound ZO1 in the membrane | $\nu_m$ | $7.5^2\pi[nm^2]$ | $7.5^2\pi/l^2$ |
| Molecular volume of solvent in the bulk | $\nu_s$ | $\frac{4\pi}{3}(7.5)^3[nm^3]$ | $\frac{4\pi}{3}(7.5)^3/l^3$ |
| Molecular volume of scaffold in the bulk | $\nu_b$ | $\frac{4\pi}{3}(7.5)^3[nm^3]$ | $\frac{4\pi}{3}(7.5)^3/l^3$ |
| **Interaction parameter** | | | |
| between free receptor and bound ZO1 in the membrane | $\chi_{mr}$ | 0 $[k_BT]$ | 0 |
| between free receptor and free lipid | $\chi_{rl}$ | 0 $[k_BT]$ | 0 |
| between ZO1 and solvent in the bulk | $\chi_b$ | 0 $[k_BT]$ | 0 |
| **Interaction parameter between bulk and membrane** | | | |
| between free lipid in the membrane and ZO1 in the bulk | $\chi_{bl}$ | 0 $[k_BT]$ | 0 |
| between free receptor in the membrane and ZO1 in the bulk | $\chi_{br}$ | $0[k_BT]$ [a] | 0 |
| between bound ZO1 in the membrane and ZO1 in the bulk | $\chi_{bm}$ | 0 $[k_BT]$ | 0 |
| **Conformational coefficient** | | | |
| Gradient coefficient of bound ZO1 in the membrane | $\kappa_m$ | $2 \times 10^{-5}$ $[\mu m^{-1}]$ | 2 $\times 10^{-4}$ |
| Gradient coefficient of free receptor in the membrane | $\kappa_r$ | $2 \times 10^{-5}$ $[\mu m^{-1}]$ | 2 $\times 10^{-4}$ |
| Gradient coefficient of scaffolds in the bulk | $\kappa$ | $2 \times 10^{-5}$ $[\mu m^{-1}]$ | 2 $\times 10^{-4}$ |
| **Mobility coefficients** | | | |
| Diffusion coefficient of scaffold in the bulk | $\Lambda_b^0 k_BT$ | 0.01 $[\mu m^2 s^{-1}]$ | 0.01 |
| Diffusion coefficient of bound scaffold in the membrane | $\Lambda_m^0 k_BT$ | 0.01 $[\mu m^2 s^{-1}]$ | 0.01 |
| Diffusion coefficient of free receptor in the membrane | $\Lambda_r^0 k_BT$ | 0.01 $[\mu m^2 s^{-1}]$ | 0.01 |

[a] Free receptor attracts the ZO1 in the bulk. However, at equilibrium, the free receptor has very low concentration and thus this interaction does not make a difference. We assume $\chi_{br} = 0$ here for convenience.

TABLE VI: **Model parameter value and their dimensionless values taken from the experimental data.** We note that the molecule sizes $\nu_m$, $\nu_s$, $\nu_b$ are values for 14-mer receptor. All other parameter values are shared in all the studies.

influences the binding affinity (see Eq. (23)). The valley arises due to the binding fraction $c$ which is considerably less than 1 for the monomer receptor based on experimental data.

From the deduced parameter values attained through fitting, it can be concluded that the 14-mer receptor exhibits a more pronounced binding affinity in comparison to the monomer receptor. Moreover, the 14-mer receptor also demonstrates a superior propensity for phase separation. This enlarged tendency suggests a preference for a condensed phase state, further amplifying its differentiation from the monomer receptor. Such findings underscore the fundamental variances in behavior between complex multimeric receptors and their simpler monomeric counterparts. Based on the derived parameters from our fitting process, it is evident that the 14-mer receptor displays a stronger binding affinity in comparison to the monomer receptor.

We further calculated the phase diagram of surface phase separation shown in Fig. 2G and 2H for monomer receptor and 14-mer receptor respectively. In the phase diagram, horizontal axis depicts scaffold protein ZO1 bulk density, while the vertical axis corresponds to the total receptor amount in the membrane. We here choose the bulk ZO1 density from 0 to 500 $nM$ approximately equal to the the experimental range. The color code denotes receptor-protein complexes density in the membrane. Intriguingly, the phase diagram for the 14-mer receptor scenario (Fig. 2H) reveals the occurrence of surface phase separation, while in contrast, no such phase separation is observed for the monomer receptor scenario (Fig. 2G). More specifically, the diagrams illustrate two distinct parameter regions. The first region, I, is characterized by a uniform concentration in the membrane, while region II represents phase-separated states in the membrane, where one phase is rich in receptor-protein complexes, and the other is not. The transition line demarcating these two states is depicted by a red solid line.

These diagrams suggest that the 14-mer receptor system possesses a higher propensity for surface phase separation compared to its monomer counterpart. This could be indicative of a difference in binding affinity or interaction strength between the two systems. As such, the 14-mer receptors might interact more robustly with the ZO1 proteins, thereby leading to an accumulation of receptor-protein complexes and subsequent phase separation. Nevertheless, without a clear understanding of the interaction parameter in the monomer receptor case, definitive predictions about the phase separation propensity of the two receptor types remain a challenge. Further experiments and computations that could reduce the uncertainty surrounding the interaction parameter would allow for a more detailed comparative study.

#### G. Derivation of dilute binding affinity given in Eq. (23)

At equilibrium, we have relation

$$\mu + q\,\mu_r = \mu_m\,. \tag{25}$$

We assume that the system is dilute, which means that $\phi_m$, $\phi_r$ and $\phi$ are non-zero but small. Using the Flory-Huggins free energy (6), (7), (8), we obtain

$$\mu = k_B T\left((1+\ln\phi) - n_b + n_b\,\chi_b + n_b\,\omega_b\right) \tag{26a}$$

$$\mu_m = k_B T\left((1+\ln\phi_m) - \ln(1-\phi_r^0) - n_m + n_m\,\chi\,(1-\phi_r^0) + n_m\,\omega\right) \tag{26b}$$

$$\mu_r = k_B T\left((1+\ln\phi_r^0) - \ln(1-\phi_r^0) - n_r + n_r\,\chi_{rl}\,(1-2\phi_r^0) + n_r\,\omega_r\right)\,. \tag{26c}$$

Combining Eqn. (25) and Eqn. (26), we get

$$\ln\phi + q\,\ln\phi_r^0 = \ln\phi_m - \Omega\,, \tag{27}$$

or equivalently,

$$\phi_m = (\phi_r^0)^q\,\phi\,e^{\Omega}\,, \tag{28}$$

where $\Omega = q - (-n_m + n_m\,\chi(1-\phi_r^0) + n_m\,\omega) + q\,(-n_r + n_r\,\chi_{rl}(1-2\phi_r^0) + n_r\,\omega_r) + (-n_b + n_b\,\chi_b + n_b\,\omega_b)$ in a constant. Here, $\Omega$ is the dilute binding affinity as it corresponds to the net affinity of binding in the dilute limit (in units of $k_B T$). In our work, we assume $\chi_{rl} = 0$, $\omega_b = \omega_r = 0$, $q = 1$, and $n_b = 1$, thus we obtain Eq. (23).

#### H. Kinetics of membrane phase separation

We also scrutinize our model by comparing the the kinetics of membrane phase separation in the experiments with the numerical simulations. To this end, we compare the spatial distribution of dense protein-rich membrane domains at specific time points. To obtain numerical results for membrane phase separation kinetics, we set the diffusivity coefficient value as $D_i = 0.01\,\mu m^2 s^{-1}$, $i = b, m, r$. The gradient coefficients $\kappa = \kappa_m = \kappa_r = 2\times10^{-5}\mu m^{-1}$. The corresponding non-dimensional values are given in Table VI. With such choices, we obtain the consistent distributions of receptor-protein-rich domains compared to the experimental patterns (see Fig. 2C, 2E in main text) at similar time scale, around 15 minutes.

Our kinetic model allows us to explore the dynamics in the bulk-membrane coupled systems with protein binding/unbinding events, facilitating an enhanced understanding of how such kinetics shape biological functions. Its ability to incorporate specific biological parameters and provide dynamic analysis makes it an invaluable tool for hypothesis testing, predictions, and potentially guiding the design of new experiments or treatments.

---

[1] Brett M. Collins, Liza Cubeddu, Nishen Naidoo, Stephen J. Harrop, Geoff D. Kornfeld, Ian W. Dawes, Paul M.G. Curmi, and Bridget C. Mabbutt. Homomeric ring assemblies of eukaryotic sm proteins have affinity for both rna and dna: Crystal structure of an oligomeric complex of yeast smf*. *Journal of Biological Chemistry*, 278(19):17291–17298, 2003.

[2] Simon Erlendsson, Thor Seneca Thorsen, Georges Vauquelin, Ina Ammendrup-Johnsen, Volker Wirth, Karen L Martinez, Kaare Teilum, Ulrik Gether, and Kenneth Lindegaard Madsen. Mechanisms of pdz domain scaffold assembly illuminated by use of supported cell membrane sheets. *eLife*, 8:e39180, jan 2019.

[3] Mason Rouches, Sarah L. Veatch, and Benjamin B. Machta. Surface densities prewet a near-critical membrane. *Proceedings of the National Academy of Sciences*, 118(40):e2103401118, 2021.

[4] Jonas Wilhelm, Stefanie Kühn, Miroslaw Tarnawski, Guillaume Gotthard, Jana Tünnermann, Timo Tänzer, Julie Karpenko, Nicole Mertes, Lin Xue, Ulrike Uhrig, Jochen Reinstein, Julien Hiblot, and Kai Johnsson. Kinetic and structural characterization of the self-labeling protein tags halotag7, snap-tag, and clip-tag. *Biochemistry*, 60(33):2560–2575, 2021.

[5] Xueping Zhao, Giacomo Bartolucci, Alf Honigmann, Frank Jülicher, and Christoph A Weber. Thermodynamics of wetting, prewetting and surface phase transitions with surface binding. *New Journal of Physics*, 23:123003, 2021.
